# Structure and conformational cycle of a bacteriophage-encoded chaperonin

**DOI:** 10.1101/2020.02.24.962803

**Authors:** Andreas Bracher, Simanta S. Paul, Huping Wang, Nadine Wischnewski, F. Ulrich Hartl, Manajit Hayer-Hartl

**Affiliations:** Department of Cellular Biochemistry, Max-Planck-Institute of Biochemistry, Martinsried, Germany

## Abstract

Chaperonins are ubiquitous molecular chaperones found in all domains of life. They form ring-shaped complexes that assist in the folding of substrate proteins in an ATP-dependent reaction cycle. Key to the folding cycle is the transient encapsulation of substrate proteins by the chaperonin. Here we present a structural and functional characterization of the chaperonin gp146 (ɸEL) from the phage EL of *Pseudomonas aeruginosa*. ɸEL, an evolutionary distant homolog of bacterial GroEL, is active in ATP hydrolysis and prevents the aggregation of denatured protein in a nucleotide-dependent manner. However, ɸEL failed to refold the encapsulation-dependent model substrate rhodanese and did not interact with *E. coli* GroES, the lid-shaped co-chaperone of GroEL. ɸEL forms tetradecameric double-ring complexes, which dissociate into single rings in the presence of ATP. Crystal structures of ɸEL (at 3.54 and 4.03 Å) in presence of ATP•BeF_x_ revealed two distinct single-ring conformational states, both with open access to the ring cavity. One state showed uniform ATP-bound subunit conformations (symmetric state), whereas the second combined distinct ATP- and ADP-bound subunit conformations (asymmetric state). Cryo-electron microscopy of apo-ɸEL revealed a double-ring structure composed of rings in the asymmetric state (3.45 Å resolution). We propose that the phage chaperonin undergoes nucleotide-dependent conformational switching between double- and single rings and functions in aggregation prevention without substrate protein encapsulation. Thus, ɸEL may represent an evolutionary more ancient chaperonin prior to acquisition of the encapsulation mechanism.

## Introduction

Chaperonins are large double-ring complexes that mediate protein folding in an ATP-dependent mechanism in all domains of life [1]. Two major groups of chaperonins exist, which transiently encapsulate non-native substrate protein (SP) for folding to proceed in an aggregation-free environment: Group I chaperonins occur in eubacteria and organelles of prokaryotic origin, mitochondria and chloroplasts (GroEL, Hsp60 and Cpn60, respectively). These chaperonins have 7-membered rings. The group II chaperonins in the cytosol of archaea and eukaryotes (thermosome and TRiC/CCT, respectively) have 8-mer rings. The subunits of both groups share a conserved three-domain architecture composed of an equatorial domain harboring the ATP binding site, an intermediate domain, which communicates nucleotide-dependent conformational changes, and a flexible apical domain for SP interaction. The equatorial domains mediate most of the inter-subunit contacts within and between the rings, whereas the apical domains form the ring opening. The group I chaperonins function together with a heptameric lid-shaped co-chaperone (GroES in bacteria, Hsp10 in mitochondria and Cpn10/20 in chloroplasts), which binds to the ends of the chaperonin cylinder and closes the folding chamber. The paradigm for the group I chaperonin mechanism is the GroEL/ES system of *Escherichia coli* [2]. GroEL and Hsp60 may transiently dissociate into single rings during their functional cycle [3, 4]. The group II chaperonins function independently of GroES-like co-chaperones and instead use helical protrusions of their apical domains as a built-in, iris-like closing mechanism [5]. Group I and II chaperonins also differ in their ring-ring contacts, with group I exhibiting staggered interactions, i.e. each subunit in one ring interacts with two subunits in the opposing ring, whereas the subunits in group II chaperonins interact in a one to one fashion.

Evolutionary more distant chaperonin homologs were discovered in the genomes of bacteriophages. While many phages encode only a GroES homolog that cooperates with the host GroEL [6], some have both GroEL and GroES homologs and few encode only a GroEL-like chaperonin [7]. The latter group includes the protein gp146 from the bacteriophage EL of *Pseudomonas aeruginosa* [8, 9], henceforth referred to as ɸEL. ɸEL shares only 22 % sequence identity with GroEL, equivalent to the evolutionary distance between group I and II chaperonins. Similar to group I chaperonins, ɸEL was reported to form a double-ring complex of 7-mer rings [9]. ɸEL was found to be expressed as early as 15 min after bacterial infection by phage, and was found to be associated with another phage protein of 32 kDa, gp188. This cell wall endolysin is unstable at 37 °C [9]. ɸEL prevented gp188 aggregation and maintained the protein in a functional state independent of host GroES [9–11]. A low resolution cryo-electron microscopy (cryo-EM) structure reported a spherical heptamer for ADP-bound ɸEL, which was suggested to represent the SP-encapsulating state [10].

Here, we performed a functional and structural characterization of the chaperonin from the phage EL of *P. aeruginosa* to better understand the ɸEL mechanism. Our results show that ɸEL functions in preventing protein aggregation but is unable to mediate the folding of rhodanese (Rho), an encapsulation-dependent model SP. In the absence of nucleotide, ɸEL forms mainly double-ring complexes, whereas single-rings are populated in the presence of ATP and physiological salt concentration. Our structural analysis by crystallography and cryo-EM failed to provide evidence of the previously reported spherical heptamer structure. The domain structures of ɸEL differ from those of GroEL by numerous insertions and deletions, which alter the putative substrate-binding cleft and the contacts at the ring-ring interface. Structural elements that might act as a built-in lid were not present in ɸEL, consistent with an evolutionary more ancient chaperone mechanism independent of substrate encapsulation.

## Materials and Methods

### Materials

Chemicals, enzymes and reagents were purchased from Merck unless otherwise noted.

### Protein production

All protein purification steps were performed at 4 °C. Protein concentrations in the final preparations were determined by measurement of absorbance at 280 nm. Purified protein samples were concentrated by ultrafiltration and snap-frozen in liquid nitrogen for storage at –80 °C.

Gene product 146 (gp146 also known as ɸEL) from *Pseudomonas* bacteriophage EL encoded by the plasmid pET22b-phi-GroEL was expressed in *E. coli* BL21(DE3) cells. Cells were grown in LB medium containing 100 mg L^-1^ ampicillin at 37 °C to an optical density of 0.5 at 595 nm. Expression of ɸEL was induced with 1 mM isopropyl β-D-1-thiogalactopyranoside (IPTG) for 3 h at 30 °C. Cells were harvested and re-suspended in ice-cold buffer PA (50 mM Tris-HCl pH 7.5). The cells were lysed by the French press method on ice using an Emulsiflex C5 apparatus (Avestin, Ottawa, Canada). After removal of cell debris by centrifugation at 120,000 g for 45 min, the supernatant was applied to a 50 ml Source 30Q column (GE Healthcare) equilibrated in buffer PA. The column was washed with 3 column volumes (CV) buffer PA containing 50 mM NaCl, followed by a linear salt gradient in buffer PA (50–500 mM NaCl, 10 CV). The eluate fractions were analyzed by SDS-PAGE. Fractions containing ɸEL (eluting at 17–21.5 mS cm^-1^) were merged and transferred into buffer PB (50 mM Tris-HCl pH 7.5, 1 mM EDTA, 1 mM DTT). Next, the material was passed over a 20 ml Heparin Sepharose FastFlow column (GE Healthcare), which was eluted with a gradient from 0 to 500 mM NaCl in buffer PB (10 CV). Finally, the concentrated fractions containing ɸEL (eluting at 8–15 mS cm^-1^) were subjected to size exclusion chromatography (SEC) on Sephacryl S-400 (GE Healthcare) in buffer, 20 mM MOPS-NaOH pH 7.2, 100 mM NaCl and 10 % glycerol.

### Size-exclusion chromatography coupled to multi-angle static light scattering (SEC-MALS)

Purified ɸEL at 2 g L^-1^ was analyzed using static and dynamic light scattering by auto-injection of the sample onto a SEC column (5 μm, 4.6×300 mm column, Wyatt Technology, product # WTC-030N5) at a flow rate of 0.35 ml min^-1^ in buffers EM50 (20 mM HEPES-NaOH pH 7.5, 50 mM KCl, 4 mM Mg acetate) or EM100 (20 mM HEPES-NaOH pH 7.5, 100 mM KCl, 2 mM Mg acetate) at 25 °C in the presence or absence of nucleotide (2 mM). The column was in line with the following detectors: a variable UV absorbance detector set at 280 nm (Agilent 1100 series), the DAWN EOS MALS detector (Wyatt Technology, 690 nm laser) and the Optilab rEXTM refractive index detector (Wyatt Technology, 690 nm laser) [12]. Molecular masses were calculated using the ASTRA software (Wyatt Technology) with the dn/dc value set to 0.185 ml g^-1^. Bovine serum albumin (Thermo) was used as the calibration standard.

### ATP hydrolysis

The ATPase activity of ɸEL (200 nM tetradecamer) at different ATP concentrations was detected by absorbance at 340 nm wavelength in low salt (LS) buffer, 50 mM Tris-HCl pH 7.5, 10 mM KCl and 10 mM MgCl_2_, or high salt (HS) buffer, 50 mM Tris-HCl pH 7.5, 100 mM KCl and 10 mM MgCl_2_, using a NADH-coupled enzymatic assay (1 mM phosphoenolpyruvate, 20 U ml pyruvate kinase, 30 U/ml lactate dehydrogenase and 0.25 mM NADH) [13]. ATPase activity assays of ɸEL and GroEL (0.2 µM tetradecamer) in the absence or presence of GroES (0.4 µM heptamer) or 0.8 µM denatured DM-MBP (diluted from 6M GuHCl, final ∼20 mM) were performed in LS or HS buffer and in presence of 1 mM ATP. The ATPase activity of ɸEL (0.2 µM tetradecamer) was also measured with increasing concentrations of GroES. The assay temperature was 25 °C.

### Rhodanese prevention of aggregation and refolding

Rhodanese (Rho; 150 μM) was denatured in 6M guanidinium-HCl (GuHCl)/10 mM DTT for 60 min at 25 °C and diluted 300-fold into buffer AP (20 mM MOPS-KOH pH 7.4, 20 mM KCl, and 5 mM MgCl_2_) containing 1 mM nucleotide in the absence or presence of ɸEL or GroEL (0.5 μM tetradecamer). Aggregation was monitored by measuring turbidity at 320 nm.

Rho refolding assays were performed as described previously with minor modifications [14]. GuHCl-denatured Rho was diluted 200-fold to a final concentration of 0.5 μM into buffer, 20 mM Tris-HCl pH 7.5, 100 mM KCl and 5 mM MgCl_2_, either lacking chaperone or containing ɸEL, ɸEL/GroES, GroEL or GroEL/GroES. The concentrations of GroEL and ɸEL were 1 μM (tetradecamer) and GroES 2 μM (heptamer). Refolding was initiated upon addition of ATP (5 mM). When indicated, chaperonin action was stopped by CDTA (10 mM). Enzymatic assays of Rho were performed as previously described [14]. Spontaneous refolding of Rho was inefficient (< 10 % yield) due to aggregation.

### Crystallization

ɸEL at 19.1 g L^-1^ in buffer, 20 mM MOPS-NaOH pH 7.3, 100 mM KCl and 2 mM Mg-acetate, was crystallized by the sitting-drop vapor diffusion method by mixing 100 nl sample with 100 nl reservoir solution using the robotics setup at the Crystallization Facility of the Max Planck Institute of Biochemistry. The drops were equilibrated against 150 μl reservoir solution at 16 °C. Crystals of ɸEL were obtained with the Complex crystallization screen [15, 16].

Crystal form I was obtained in presence of 2 mM ATP, 5 mM BeSO_4_ and 20 mM NaF with a reservoir solution containing 5 % PEG-4000, 0.2 M Na-acetate and 0.1 M Na_3_-citrate pH 5.5.

Crystal form II was obtained in presence of 2 mM ATP, 5 mM BeSO_4_ and 20 mM NaF with a reservoir solution containing 8 % PEG-6000, 0.15 M NaCl and 0.1 M Tris-HCl pH 8.0.

For vitrification, the crystals were sequentially incubated in reservoir solution containing additionally 12.5 and 25 % glycerol for 15 min each and were then rapidly cooled in liquid nitrogen.

### Diffraction data collection and processing

The X-ray diffraction data were collected by the oscillation method at beamline ID30B of the European Synchrotron Radiation Facility (ESRF) in Grenoble, France. All data were integrated and scaled with XDS [17]. Pointless [18], Aimless [19] and Ctruncate [20], as implemented in the CCP4i graphical user interface (GUI) [21], were used for data reduction.

### Crystal structure solution and refinement

The space group symmetry and size of the asymmetric unit suggested that crystal forms I and II contained single-ring heptamers at 65 % solvent content, which is within the expected range for protein crystals (∼75–40 % solvent) [22]. Analysis of the self-rotation function calculated with Molrep [23] indicated the presence of seven-fold non-crystallographic symmetry (NCS) consistent with the presence of single-ring heptamers in the crystal lattice. For solving the structure of crystal form II by molecular replacement, the cryo-EM density for the heptadecameric ɸEL•ATP at 6.8 Å resolution (EMDB entry EMD-6492, [10]) was segmented with Chimera [24] and the density for a single-ring heptamer extracted. With this density as a search model, a plausible molecular replacement solution was obtained. The density modification program Resolve [25] was used to extend the phases beyond 6.8 Å employing seven-fold NCS averaging and refinement. After B-factor sharpening, the resulting experimental electron density revealed features of secondary structure sufficient for manual model building with Coot [26]. ɸEL crystal form I was solved by molecular replacement using Molrep [23] with these coordinates as a search model. To identify the nucleotides bound to subunits of ɸEL, omit maps were calculated after applying random coordinate shifts to the preliminary model with Pdbset (to suppress model bias) and refinement with Refmac5 [27], which revealed density consistent with ADP-bound to chains B, D and F in crystal form II, and with ATP/ADP•BeF_x_ in all other chains. Since ATP and ADP•BeF_x_ cannot be distinguished at the resolution, and since ATP was added to the crystallization mix, ATP was included into the model. The models were refined with Refmac5 using local NCS restraints and translation-libration-screw (TLS) parametrization of B-factors [27]. Residues 1 and 553–558 were disordered in all chains of the final models. Furthermore, no interpretable density was observed for loop residues 290–294 in crystal form I and in chains A, C, E and G of crystal form II. Residues facing solvent channels with disordered sidechains were truncated after C-β.

Crystallographic structure factors and model coordinates have been deposited to wwPDB under accession numbers 6TMT and 6TMU.

### Cryo-electron microscopy and single particle analysis

The samples were prepared by mixing equal volumes of ɸEL stock solution (2.25 g l^-1^) in buffer, 20 mM HEPES-NaOH pH 7.5, 100 mM NaCl and 1 mM EDTA, and dilution buffer (20 mM HEPES-NaOH pH 7.5 and 8 mM MgCl_2_) containing either no nucleotide or 4 mM ADP or ATPγS. The dilution buffer with ADP also contained 0.08 % n-octyl-β-D-glucopyranoside. Subsequently the mixture was incubated at room temperature for 5 min. Holey carbon supported copper-grids (Quantifoil R2/1 300 mesh) were plasma-cleaned for 30 s (Harrick Plasma) immediately before use. All cryo-grids were prepared using a Vitrobot Mark 4 (FEI) by applying sample (5 μl) to a plasma-cleaned grid at 25 °C and 100 % humidity, then semi-automatically blotted for 2 s and plunge-frozen in liquid ethane.

The grids with ɸEL•ADP or ɸEL•ATPγS were analyzed on a Talos Arctica (FEI) transmission electron microscope (TEM) at 200 kV. Frames were recorded with a Falcon 3EC direct detector (FEI) operated in movie mode at 0.05 s per frame, at a pixel size of 1.997 Å and 2.019 s total exposure, with an estimated cumulative dose of 42–43 e^-^ Å^-2^. EPU (FEI) software was used for automated data collection. MotionCor2 [28] was employed to correct the movie stacks for beam-induced motion and dose weighting. Ctffind4.1 [29] was used to estimate the defocus. For generating templates used in auto-picking, 6,668 single particles of ɸEL•ADP were picked manually and subjected to 2D classification with RELION 3.0 [30, 31]. The eight largest 2D classes were selected and used in Gautomatch (http://www.mrc-lmb.cam.ac.uk/kzhang/Gautomatch) as templates for automated particle picking.

The grid with apo-ɸEL was analyzed on a Titan Krios (FEI) TEM at 300 kV with a pixel size of 1.09 Å. Data were collected with a K3 direct detector (Gatan) recording 50 frames per movie during 5.992 s total exposure with an estimated cumulative dose of 77.6 e^-^ Å^-2^. SerialEM software was used for automated data collection [32, 33]. Motion correction, dose weighting, defocus analysis and particle picking were carried out automatically during data collection using the Focus software package [34]. Movies with large drift, exhibiting ice diffraction or poor resolution (> 5 Å) in the power spectra were immediately discarded.

In absence of nucleotide and in presence of ADP, ɸEL had a tendency to associate into large aggregates. Images with thick aggregates were removed after visual inspection. RELION 3.0 was used for further data processing [31]. The complete data sets went through two rounds of 2D classification to remove contaminations or false positive particles. Using RELION 3.0, an initial model was generated from the remaining particles and used as a reference map for symmetry-free 3D refinement. The aligned particles were subjected to 3D classification. The particles from the largest class were used for further 3D refinement and post-processing including mask application and B-factor sharpening. The resulting electron density maps were inspected with Chimera [24], and *C*2 (apo-ɸEL) and *D*7 (ɸEL•ADP and ɸEL•ATPγS) particle symmetry were detected. Re-processing of the particles using the same protocol as above, but with application of symmetry restraints, yielded improved maps for manual model building with Coot [26]. First, the apical, intermediate and equatorial domains of ɸEL from the form-I crystal structure were separately placed into density by rigid-body real-space fitting. After manual adjustment of the coordinates to the density, the models were refined with Refmac5 in reciprocal space, using jelly-body and NCS restraints [27].

Electron density maps for apo-ɸEL, ɸEL•ADP and ɸEL•ATPγS have been deposited to EMDB under accession numbers 10528, 10529 and 10530, respectively. The corresponding model coordinates have been deposited to wwPDB under accession codes 6TMV, 6TMW and 6TMZ.

### Structure analysis

The quality of the structural models was analyzed with the program Molprobity [35]. Coordinates were aligned with Lsqkab and Lsqman [36]. Figures were generated with the programs Pymol (http://www.pymol.org), Chimera [24] and ESPript [37].

## Results

### Oligomeric state of ɸEL

ɸEL was expressed as a soluble protein in *E. coli* and purified to apparent homogeneity (S1A Fig). To determine the oligomeric state of ɸEL at different ionic strength and in the absence or presence of nucleotide (at 25 °C), we subjected ɸEL to size exclusion chromatography combined with multi-angle light scattering (SEC-MALS). At 100 mM KCl, ɸEL fractionated at a molecular weight (MW) of ∼770 kDa (Fig 1A), indicating a high population of double-ring complexes (theoretical MW ∼863 kDa). A similar MW (∼725 kDa) was observed in the presence of ADP (Fig 1B). In contrast, ɸEL in the presence of ATP shifted to a homogeneous population of complexes with a MW of ∼409 kDa, close to the MW of the single-ring heptamer (theoretical MW ∼431 kDa) (Fig 1C). This suggests that binding of ATP to ɸEL in the presence of close to physiological salt concentration destabilizes the inter-ring contacts in the double ring structure. However, at a lower salt concentration of 50 mM KCl, this destabilization was not observed and mainly double-ring complexes were populated, as for apo and ADP-bound ɸEL (Figs 1A-C). These results suggests that under conditions of ongoing ATP-binding and hydrolysis (at 100 mM KCl), ɸEL may cycle between double-ring and single-ring complexes.

**Fig 1.**
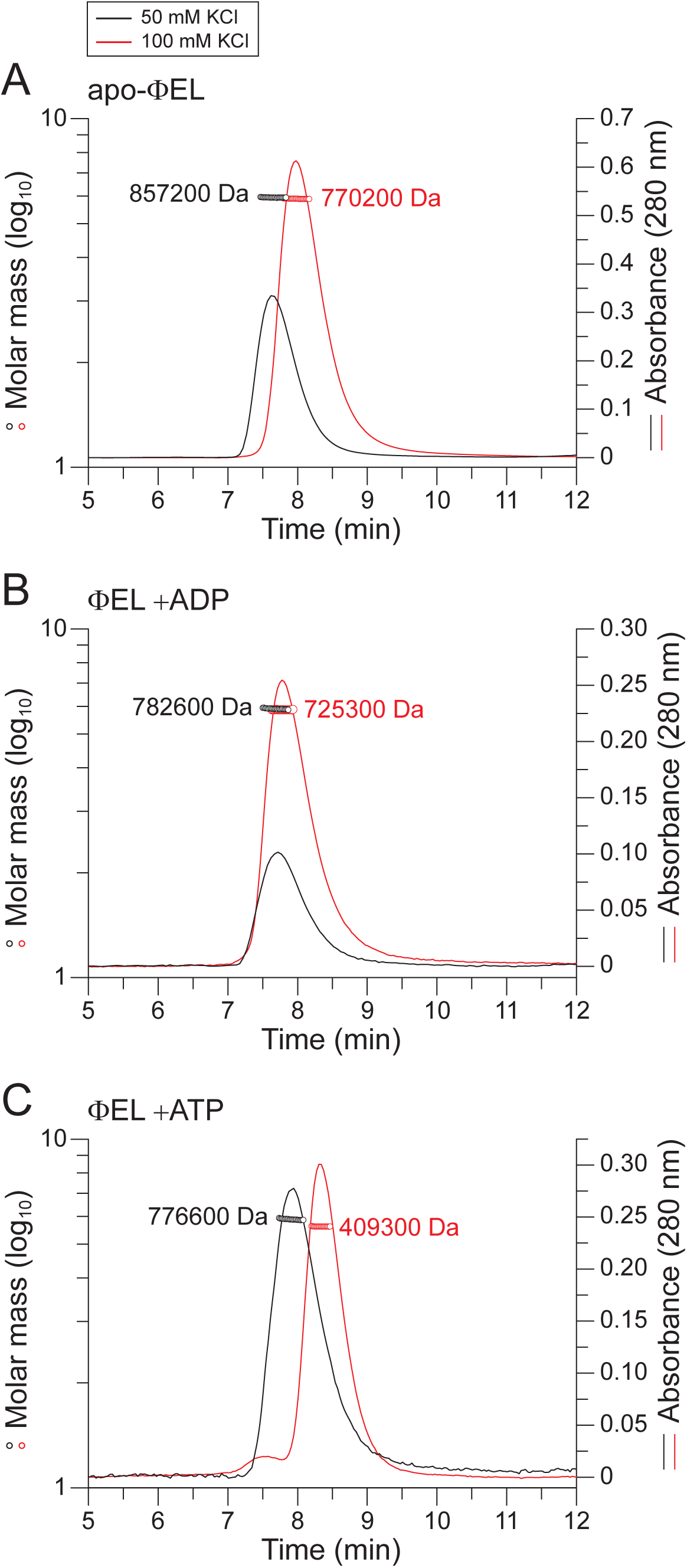
Oligomeric state of ɸEL in solution. (A-C) SEC-MALS analysis of ΦEL in absence of nucleotide (A), in presence of 2 mM ADP (B) or 2 mM ATP (C). The chromatographic absorbance traces at 280 nm wavelength are shown. The molecular mass determined for the protein peaks by static light scattering is indicated. The black and red traces were recorded in presence of 50 and 100 mM salt, respectively.

### ATPase activity of ɸEL

ATP hydrolysis by group I and II chaperonins exhibits positive cooperativity within rings, with higher ATP occupancy triggering ATP hydrolysis (Hill coefficient for GroEL ∼2.8) [38]. In addition, negative cooperativity between the rings results in a reduced hydrolysis activity at still higher ATP concentration [38]. At 25 °C and 10 mM KCl, ɸEL hydrolyzed ATP with near Michaelis-Menten kinetics, reaching a maximal turnover number of 558 ± 28 ATP min^-1^ per tetradecamer, which is ∼8-fold higher than that of *E. coli* GroEL [4, 38]. A Hill coefficient of 1.21 ± 0.05 and an apparent K_M_ of 0.68 ± 0.07 mM ATP were determined (Fig 2A). At 100 mM KCl, we measured a similar maximal turnover number (576 ± 7 ATP min^-1^ per tetradecamer), a Hill coefficient of 1.61 ± 0.18 and a ∼10-fold lower K_M_ of 0.072 ± 0.008 mM ATP (Fig 2B). Thus, at the near physiological salt concentration the affinity of ɸEL for ATP is increased with weak positive cooperativity for ATP hydrolysis. No evidence for negative cooperativity in ATP hydrolysis was detected. The higher ATPase activity and lower Hill coefficient of ɸEL compared to GroEL are consistent with a reduced level of allosteric coordination within and between rings of ɸEL.

**Fig 2.**
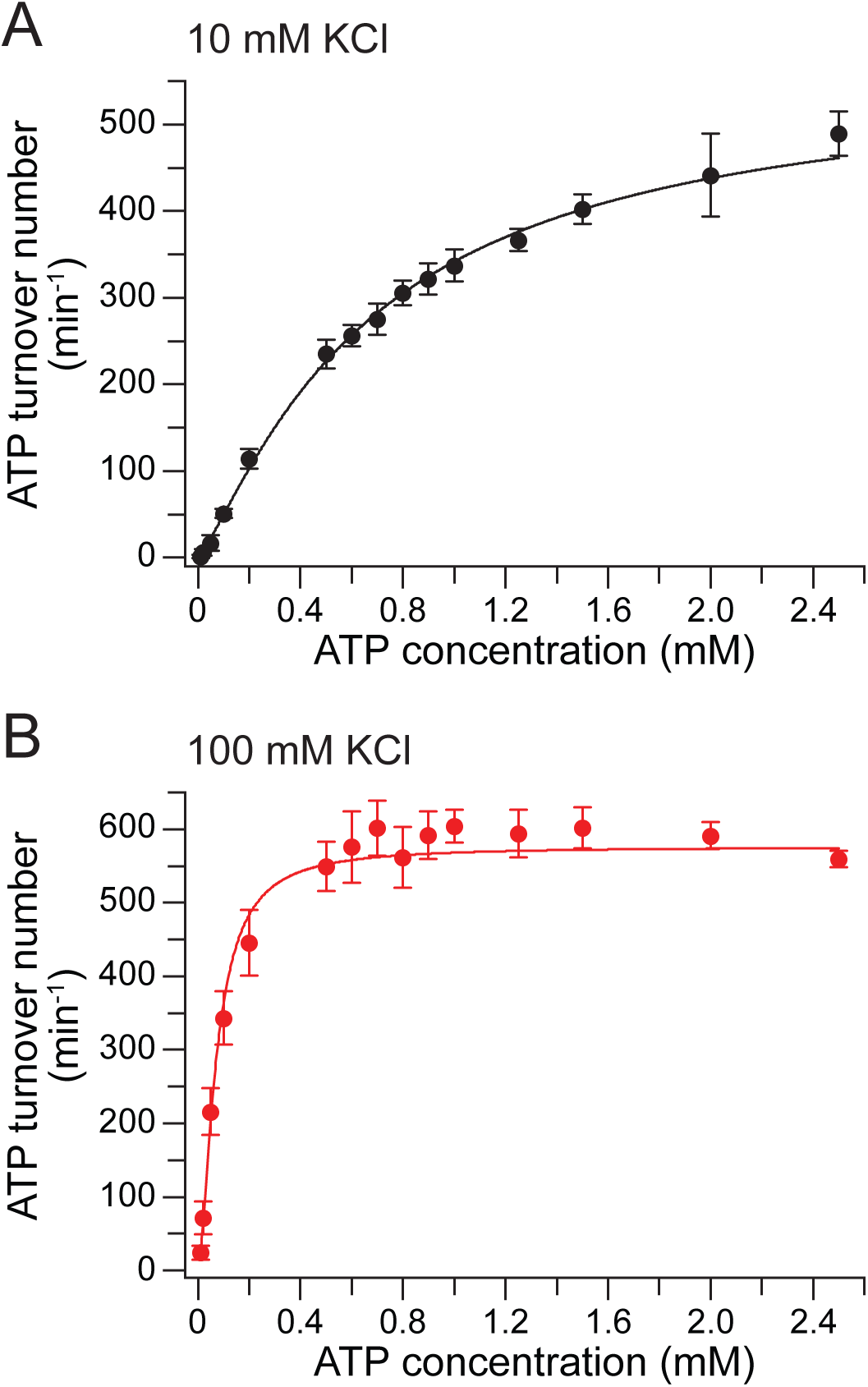
Concentration dependence of ATP hydrolysis by ɸEL. (A, B) The curves show the concentration dependence of ATP hydrolysis by ɸEL in presence of 10 (A) and 100 mM KCl (B). Shown are the averages of three experiments. Error bars indicate standard deviations. The red lines represent the Hill curve fittings of the data.

### No interaction of ɸEL with GroES

The co-chaperone GroES (2-fold molar excess) inhibited the GroEL ATPase activity by ∼50 % both at 10 and 100 mM KCl (Figs 3A and 3B) [39]. In contrast, we observed no effect of GroES on the ATPase activity of ɸEL (at 1 mM ATP) independent of salt concentration (Figs 3A and 3B). Note that at a concentration of 1 mM ATP the ATPase rate of ɸEL measured at 10 mM KCl is ∼50 % lower than in the presence of 100 mM KCl. Increasing GroES up to 8-fold excess over ɸEL also showed no effect (S1B Fig), suggesting the absence of a functional interaction. Note that *E. coli* GroES shares 61 % amino acid identity (90 % similarity) with the GroES of the host bacterium of phage EL. Specifically, the GroES mobile loop sequences that mediate binding to the apical domains of GroEL are essentially identical (82 % identity, 94 % similarity) between *E. coli* GroES and *P. aeruginosa* GroES. To further analyze a possible interaction of GroES with ɸEL, we took advantage of the fact that the mobile loops of GroES become protected against degradation by proteinase K (PK) upon complex formation with GroEL in the presence of ADP (Fig 3C, lanes 1, 2, 9 and 10) [40]. In contrast, no protection of GroES was observed in the presence of ɸEL (Fig 3C, lanes 5 and 6), suggesting absence of binding. Note that PK cleaved ɸEL into fragments of ∼45 and ∼20 kDa (Fig 3C, lanes 3 and 4), while GroEL is largely PK resistant (Fig 3C, lanes 7 and 8), except for cleavage of the flexible 16 C-terminal residues of the GroEL subunits [41].

**Fig 3.**
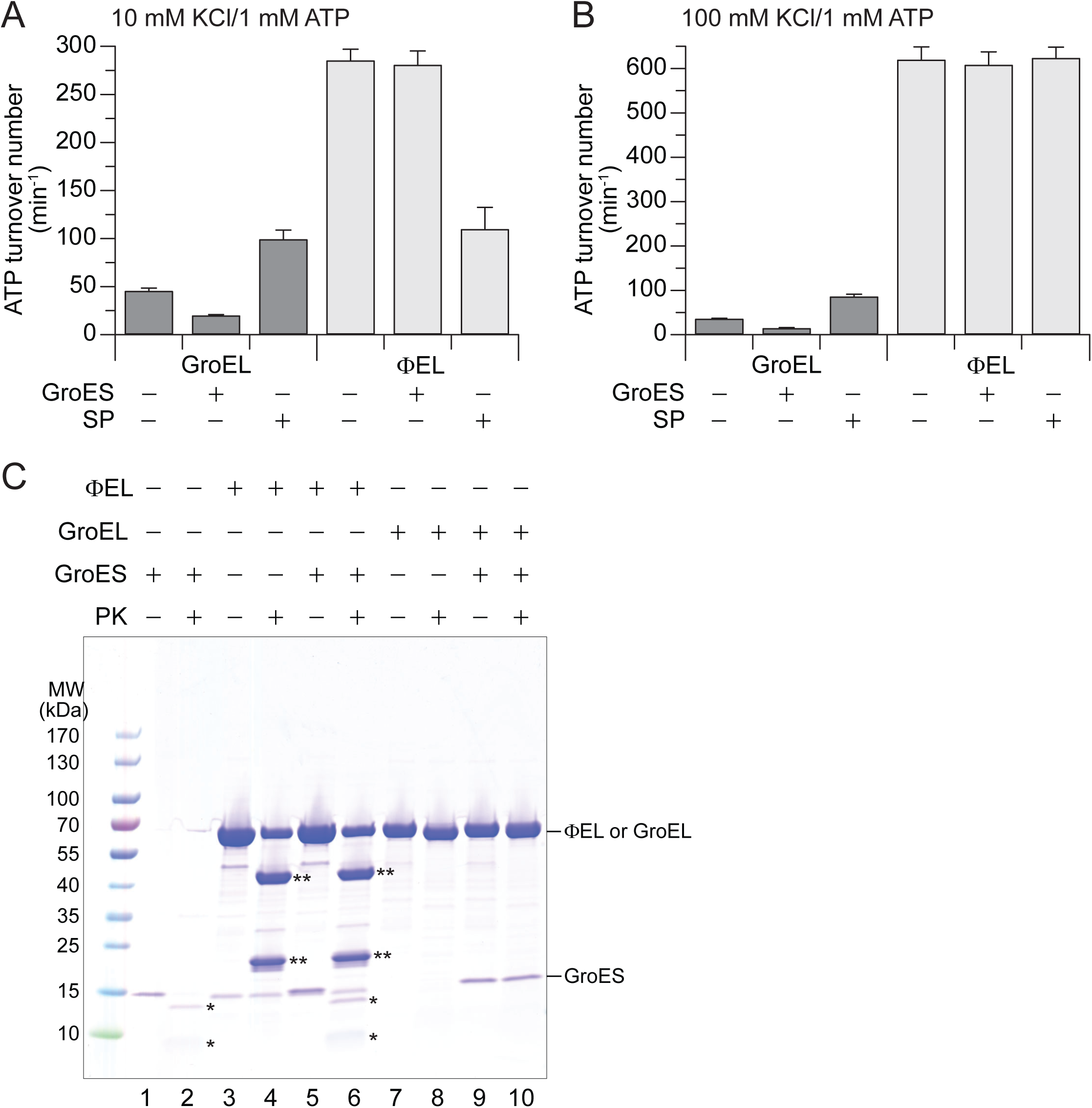
Effect of GroES on GroEL and ɸEL. (A, B) ATPase activity of GroEL and ɸEL in presence and absence of a two-fold excess of GroES or the model substrate DM-MBP. The buffer contained 10 mM (A) or 100 mM KCl (B) and the ATP concentration was 1 mM. The bar graph shows the averages from at least three experiments; error bars indicate standard deviations. (C) Proteinase K (PK) sensitivity of GroES alone, and of ɸEL and GroEL in presence and absence of GroES. The experiment was performed in buffer 20 mM Tris-HCl pH 7.5, 100 mM KCl, 5 mM MgCl_2_ and 0.2 mM ADP [42]. The concentrations of GroEL and ɸEL were 1.5 μM; GroES was at 1.0 μM. After 3 min incubation with 0.2 g L^-1^ PK at 25 °C, the protease reaction was stopped by addition of phenylmethylsulfonyl fluoride (1 mM). The mixtures were analyzed by SDS-PAGE. *, proteolytic fragments of GroES; **, proteolytic fragments of ɸEL.

We next analyzed the effect of SP on the ATPase of ɸEL. We used as model SP the double mutant of maltose binding protein (DM-MBP), which folds slowly (t_1/2_ ∼35 min at 25 °C) [43] in the absence of chaperonin and has a low aggregation propensity, thus allowing us to perform ATPase measurements under SP saturation of chaperonin [44]. Binding of non-native DM-MBP simulated the GroEL ATPase by ∼2-fold independent of salt concentration (Figs 3A and 3B). Interestingly at 10 mM KCl, DM-MBP inhibited the ɸEL ATPase by ∼60 % (Fig 3A). In contrast, no inhibition of ɸEL ATPase was observed at 100 mM KCl (Fig 3B). Thus, the effect of SP on the ATPase appears to depend on whether ɸEL populates double-ring complexes (low salt) or single-rings (high salt) (Fig 1C).

In summary, GroEL and ɸEL differ substantially with regard to their interaction with co-chaperone (*E. coli* GroES) and SP.

### Chaperone activity of ɸEL

Next, we investigated the molecular chaperone activity of ɸEL. We first tested the ability of ɸEL to prevent aggregation of the model chaperonin SP rhodanese (Rho; ∼30 kDa). This protein has a high propensity to aggregate upon dilution from denaturant into buffer, but folds efficiently upon transient encapsulation in the GroEL-GroES cage [42, 45–48]. Aggregation was monitored spectrophotometrically by measuring turbidity at 320 nm. A time-dependent aggregation of Rho was observed upon dilution from denaturant to a final concentration of 0.5 μM (Figs 4A and 4B). Note that Rho aggregation was independent of the presence of nucleotide (Figs 4A and 4B). GroEL at a 1:1 molar ratio, in the absence of nucleotide, completely prevented the aggregation of Rho in the turbidity assay. In contrast, ɸEL failed to prevent Rho aggregation (Fig 4A). However, ɸEL inhibited Rho aggregation with high efficiency in the presence of ATP or ADP (Figs 4A and 4B). In the case of GroEL, aggregation was efficiently prevented in the presence of ADP (Fig 4B), whereas addition of ATP reduced the ability of GroEL to bind non-native Rho [45] (Fig 4A). Thus, in contrast to GroEL, SP binding to ɸEL requires presence of nucleotide.

**Fig 4.**
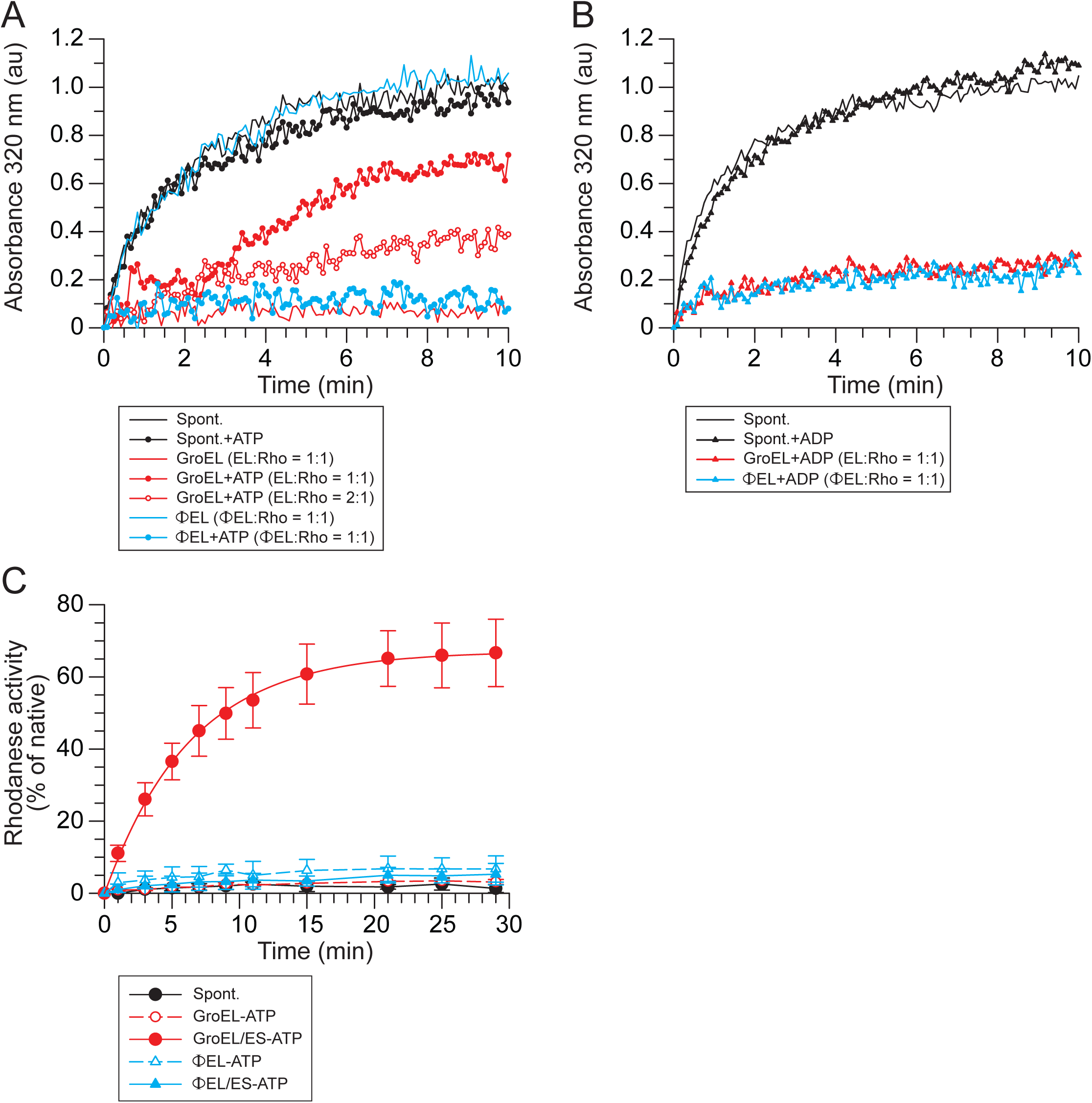
Molecular chaperone activity of ɸEL. (A, B) Aggregation prevention activity of ɸEL and GroEL in presence and absence of ATP (A) or ADP (B). Rhodanese (Rho) aggregation was monitored by turbidity assay at 320 nm wavelength. The results of representative experiments are shown. (C) Rho refolding in presence of ATP. The chaperones GroEL, GroES and ɸEL were present when indicated. After quenching ATP hydrolysis by addition of CDTA at the indicated time points, Rho enzyme activity was determined. The averages from three experiments are shown, error bars indicate standard deviations.

Rho activity assays were performed to determine whether aggregation prevention by ɸEL was coupled to productive folding. Control reactions showed only ∼5 % recovery of Rho enzyme activity during spontaneous folding (Fig 4C), due to Rho aggregation. Efficient refolding was obtained by GroEL in a manner dependent on GroES and ATP [42, 45]. In contrast, only ∼8 % of Rho enzyme activity was recovered in the presence of ɸEL and ATP (Fig 4C). Addition of GroES was without effect (Fig 4C), consistent with the absence of a functional interaction of ɸEL with GroES (Fig 3 and S1B Fig).

To investigate the fate of Rho during attempted refolding with ɸEL and ATP, we analyzed the reaction after 30 min by SEC in the presence of ATP, followed by SDS-PAGE and immunoblotting with anti-Rho. ɸEL fractionated in an asymmetric distribution (with the peak in fraction 8), consistent with cycling between double- and single-ring complexes (S2A Fig). Rho fractionated with a higher mass than ɸEL (with the peak in fraction 7), indicating that it formed a high molecular weight aggregate as it dissociated from ɸEL during gel filtration (S2A Fig). Rho refolding reactions with GroEL and GroES were analyzed as a control. In the absence of ATP, Rho co-fractionated with GroEL (S2B Fig), whereas in the presence of ATP most of the Rho fractionated as the monomeric native protein (S2C and S2D Figs).

In conclusion, ɸEL can bind non-native Rho in the presence of nucleotide thereby preventing aggregation. However, ATP-dependent cycling of Rho fails to promote productive folding, independent of the presence of GroES. Apparently, during cycling Rho is transiently released in an unfolded state, explaining its aggregation during gel filtration when rebinding to ɸEL is precluded.

### Crystal structures of ɸEL in presence of ATP•BeFx

To obtain insight into the structural features underlying the ATP-dependent double- to single-ring transition in ɸEL, we tried to determine the crystal structures of apo-ɸEL and of ɸEL in presence of ADP or ATP•BeF_X_ (ADP•BeF_X_ is a mimic of bound ATP prior to hydrolysis). However, only crystals of ɸEL with ATP•BeF_X_ diffracted below 7 Å resolution. Two crystal forms were identified that diffracted to 4.03 and 3.54 Å resolution (Table 1). The structure was solved by molecular replacement at 6.8 Å resolution using the cryo-EM map of the ɸEL-ATP tetradecamer as a search model (EMD-6492) [10]. The seven-fold non-crystallographic symmetry was then employed to extend the phases to 3.54 Å, resulting in an electron density map of sufficient quality to build an atomic model. The other crystal form was solved by molecular replacement using this model. Both structures consist of heptameric single-ring complexes of ɸEL (Figs 5A and 5B).

**Fig 5.**
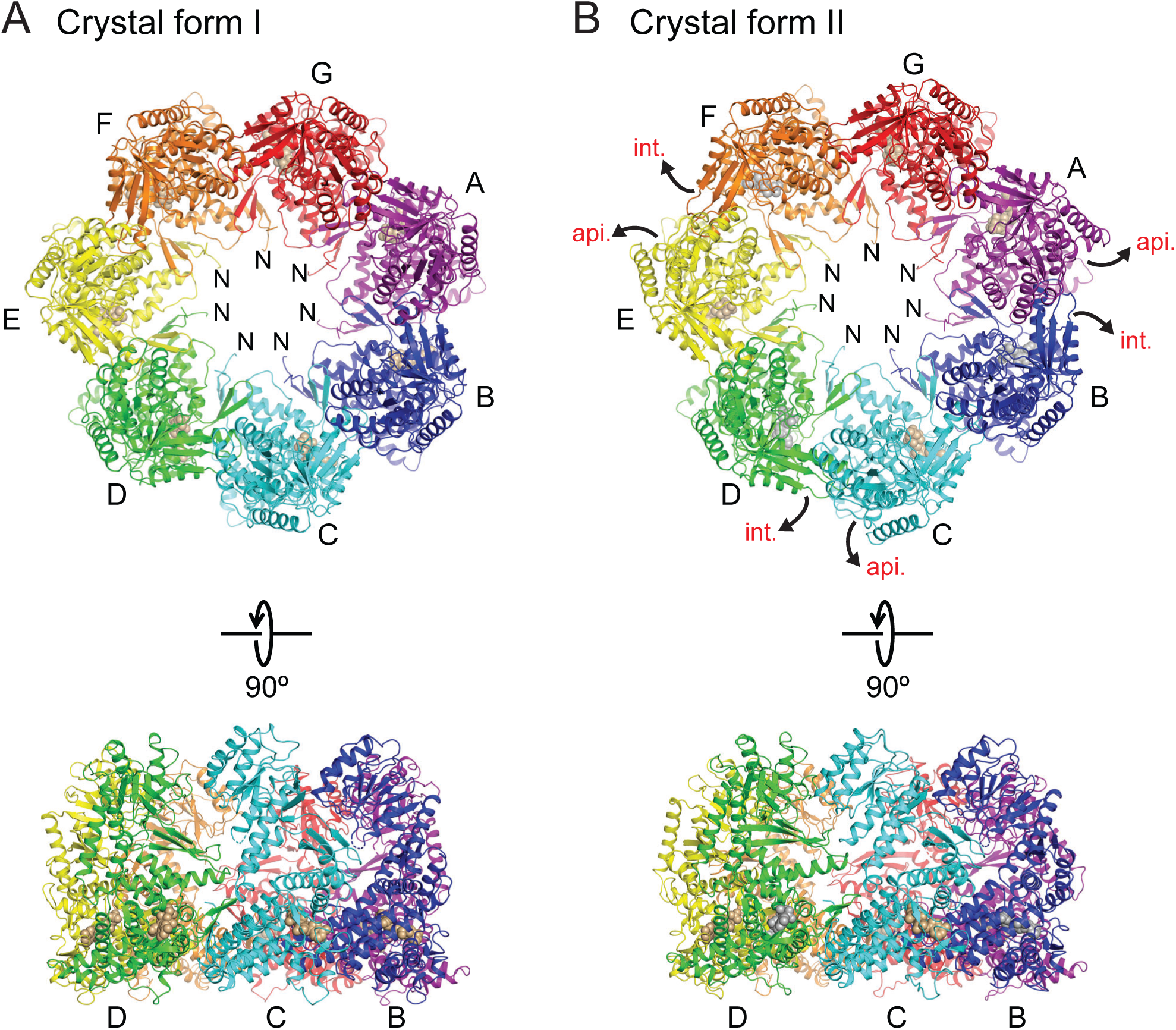
Crystal structures of ɸEL. (A) Ribbon representation of the ɸEL complex in crystal form I. Perpendicular views of the ɸEL complex are shown. The subunits are shown in rainbow colors. Bound ATP and Mg^2+^ is colored beige and shown in space-filling mode. Subunit chain identifiers and N-termini are indicated. (B) Structure of the ɸEL complex in crystal form II, using the same representation style. Bound ADP is colored silver and shown in space-filling mode. Domain movements in ɸEL subunits compared to crystal form I are indicated by curved arrows.

**Table 1.** Crystallographic data collection and refinement statistics.

| Crystal form | I | II |
| --- | --- | --- |
| PDB code | 6TMT | 6TMU |
| **Data Collection** |  |  |
| Nucleotide | ATP•BeF<sub>x</sub> | ATP•BeF<sub>x</sub> |
| Diffraction source | ESRF beamline<br>ID30B | ESRF beamline<br>ID30B |
| Space group | *P*2<sub>1</sub>2<sub>1</sub>2<sub>1</sub> | *P*2<sub>1</sub>2<sub>1</sub>2<sub>1</sub> |
| Cell dimensions<br>a, b, c (Å) | 137.9, 149.4,<br>268.8 | 145.5, 151.5,<br>261.3 |
| $\alpha, \beta, \gamma$ (°) | 90, 90, 90 | 90, 90, 90 |
| Wavelength (Å) | 0.85000 | 0.85000 |
| Resolution (Å) | 48.98 – 4.03<br>(4.17 – 4.03)* | 49.47 – 3.54<br>(3.62 – 3.54) |
| *R*<sub>merge</sub> | 0.344 (1.891) | 0.111 (1.119) |
| *R*<sub>p.i.m.</sub> | 0.099 (0.528) | 0.056 (0.557) |
| *I*/σ*I* | 6.9 (1.6) | 7.9 (1.3) |
| Completeness (%) | 99.9 (100) | 99.8 (99.9) |
| Redundancy | 12.9 (13.7) | 4.9 (4.9) |

| Refinement |  |  |
| --- | --- | --- |
| Resolution (Å) | 30 – 4.03 | 30 – 3.54 |
| No. reflections | 44298 | 67159 |
| $R_{\text{work}} / R_{\text{free}}$ | 0.2562 / 0.2991 | 0.2504 / 0.2747 |
| No. atoms |  |  |
| Protein | 29155 | 29093 |
| Nucleotides | 224 | 213 |
| Average $B$-factors | | |
| Protein (Å<sup>2</sup>) | 175 | 176 |
| Nucleotides (Å<sup>2</sup>) | 112 | 151 |
| R.m.s. deviations |  |  |
| Bond length (Å) | 0.007 | 0.005 |
| Bond angles (°) | 1.067 | 0.967 |
| Ramachandran plot |  |  |
| Favored (%) | 94.5 | 95.2 |
| Allowed (%) | 5.3 | 4.7 |
| Outliers (%) | 0.2 | 0.1 |

\* Values in parenthesis are for outer shell.

As anticipated, the ɸEL subunits displayed the typical three-domain architecture of chaperonins, composed of an equatorial domain (residues 2–130 and 425–552), an intermediate domain (residues 131–188 and 388–424), and an apical domain (residues 189–387) (Fig 6A). These domains formed rigid-body units in the heptamer rings, which tend to move *en bloc* when comparing subunits (S3A-D Figs). The apical domain appeared to be the most mobile units, as judged from their comparatively poor electron density when not stabilized by crystal contacts. The equatorial domains were well-defined, as they form the majority of the contacts between adjacent subunits, similar to known chaperonin structures.

**Fig 6.**
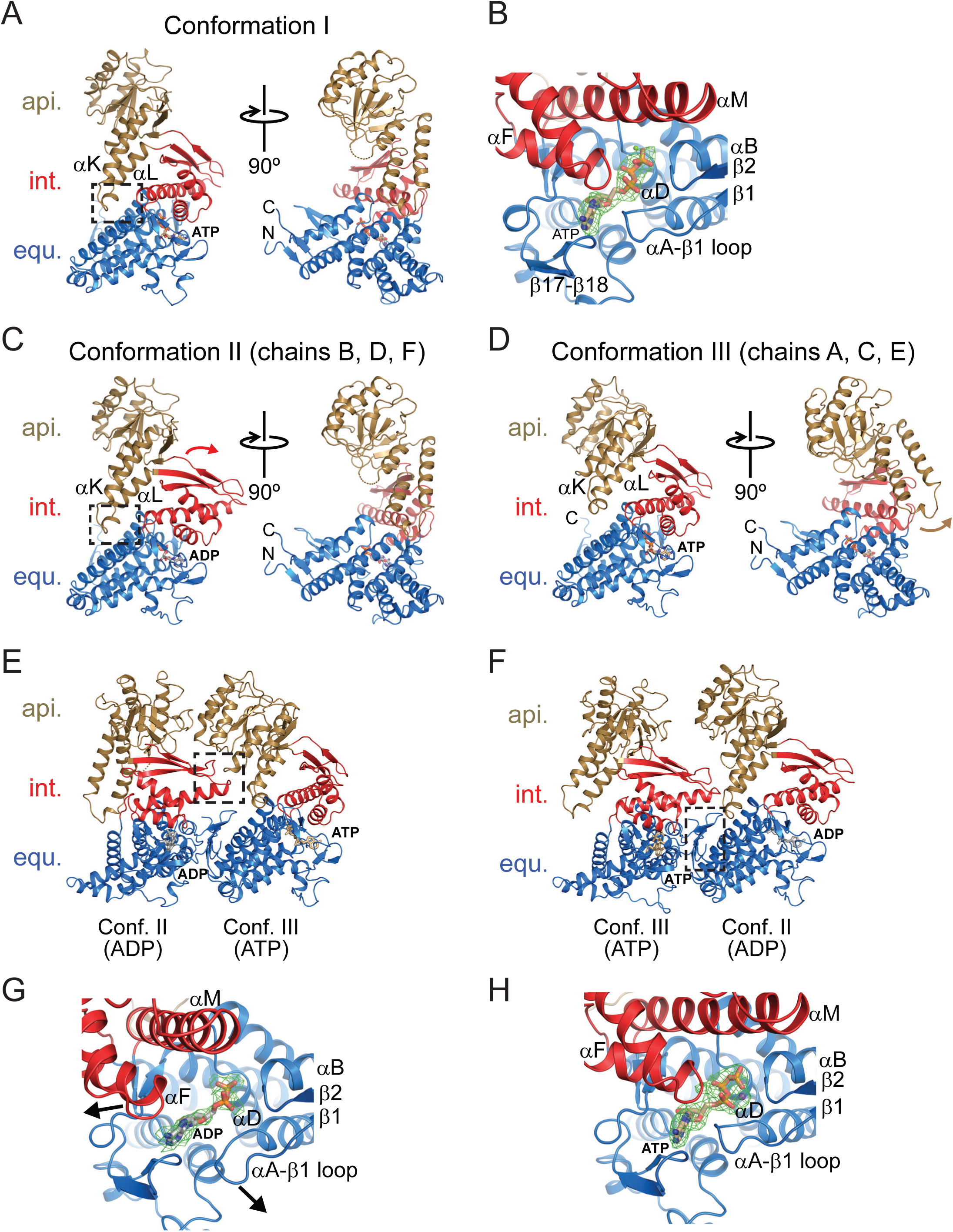
Conformations of ɸEL subunits observed in the crystal structures. (A) Ribbon representation of ɸEL subunit conformation I as observed in chain A of crystal form I. Perpendicular views of ɸEL are shown. Apical, intermediate and equatorial domain are shown in gold, red and blue, respectively. Bound ATP is shown in stick representation. The intramolecular contact between apical and equatorial domain is highlighted by a green box. The αK-αL hairpin is indicated. N- and C-termini are indicated. (B) Zoom-in on the bound ATP and Mg^2+^ in chain A of crystal form I (conformation I). The representation is equivalent to panel A. Unbiased *F*o-*F*c omit density at 2.85σ for the nucleotide is shown in green as meshwork. Secondary structure elements involved in contacts to the nucleotide are indicated. (C) Ribbon representation of ɸEL subunit conformation II as observed in chain D of crystal form II. The rotation of the intermediate domain (red) compared to conformation I is indicated by a curved arrow. Bound ADP is shown in stick representation. (D) Ribbon representation of ɸEL subunit conformation III as observed in chain A of crystal form II. The rotation of the apical domain (gold) compared to conformations I and II is indicated by a curved arrow. Bound ATP is shown in stick representation. (E) Contact between subunits in conformations II and III, as observed between chains E and D in crystal form II. The characteristic intermediate-apical domain intermolecular contact is highlighted by a box. (F) Contact between subunits in conformations III and II, as observed between chains B and A in crystal form II. The contacts are almost exclusively formed between the equatorial domains (boxed area). (G) Zoom-in on the bound ADP in chain D of crystal form II (conformation II). Unbiased *F*o-*F*c omit density at 1.7σ for the nucleotide is shown in green as meshwork. Movement of the intermediate domain and remodeling of the αA-β1 loop, respectively, are indicated by arrows. (H) Zoom-in on the bound ATP and Mg^2+^ in chain A of crystal form II (conformation III). Unbiased *F*o-*F*c omit density at 2.85σ for the nucleotide is shown in green as meshwork.

### Structure of ɸEL subunits in crystal form I

The ɸEL heptamer in crystal form I (4.03 Å resolution) consisted of subunits in closely similar conformations (conformation I, r.m.s.d. range for Cα positions 0.28–1.36 Å) (S3A Fig). In conformation I, the tip of the protruding αK-αL helical hairpin of the apical domain contacts the equatorial domain (Fig 6A). The αK-αL helical hairpin makes additional contacts to the intermediate domain (Fig 6A). These contacts appear to stabilize the orientation of the apical domain. No such contact is found in group I chaperonin structures (group II chaperonins do not contain this helical hairpin). There were no detectable contacts between apical domains unless forced by crystal packing.

All subunits as judged from the electron density had either ATP or ADP•BeF_X_ bound in the equatorial domain. For simplicity, we modelled the bound nucleotide as ATP. ATP was cradled by the αA-β2 loop (residues 30–33), the N-terminal ends of helix αD (residues 86–90) and helix αN1 (residues 428–430), helix αO (residues 474 and 478) and residues 504–506 and 519–521 (Fig 6B). Thr145 and Gln149 at the C-terminal end of αF in the intermediate domain approached the nucleobase, but did not appear to make full contact (Fig 6B). Intermediate domain helix αM with the catalytic residue Asp412 (equivalent to Asp398 in *E. coli* GroEL) was at ∼6.5 Å distance to the γ-phosphate of ATP, consistent with a conformation poised for ATP hydrolysis (Fig 6B).

### Structure of ɸEL subunits in crystal form II

In contrast to crystal form I, the subunits in crystal form II exhibited considerable conformational differences (r.m.s.d. range 0.60–4.43 Å), and two new states could be assigned (conformations II and III) (Fig 6A and S3A-D Figs). Subunits with conformation II (chains B, D, F) alternate with subunits in conformation III (chains A, C, E) around the ring, with the remaining chain G having conformation I (S3A-D Figs). Superposition of the subunits in conformations II and III showed that the apical domain can pivot by ∼22° around the joint with the intermediate domain (as determined with the program DynDom [49]) (Figs 6C and 6D; S3B and S3C Figs). Notably, in four of the subunits (chains B, D, F and G), the tip of the αK-αL helical hairpin protruding from the apical domain is oriented as in crystal form I (Fig 6C; S3B and S3D Figs). In chains A, C and E (conformation III) the apical domain is re-oriented, allowing residues 290–294 to contact the tip of the intermediate domain of the adjacent subunit (chains B, D and F; conformation II) (Figs 6D and 6E). This re-orientation of the apical domains of chains A, C and E also results in the αK-αL helical hairpin to point outwards into the solvent (Fig 6D and S3C Fig). The apical-intermediate domain intra-ring contact also requires a 19–21° outward rotation of the intermediate domains in chains B, D and F (Fig 6C and S3B Fig). All other inter-subunit contacts are limited to the equatorial domains (Fig 6F).

All subunits in crystal form II contained electron density for a bound nucleotide (Figs 6G and 6H). The orientation of the intermediate domain correlated with the identity of the bound nucleotide: The subunits with outward-oriented intermediate domain (chains B, D and F) contained weak nucleotide density, consistent with partial occupancy by ADP (Fig 6G). The β-phosphate of ADP was coordinated by the amides at the N-terminal end of helix αD; additional density for BeF_x_ was not detectable. The nucleotide electron density for the subunits with inward-oriented intermediate domain (chains A, C, E and G) was consistent with nucleoside triphosphate, modelled as ATP and Mg^2+^ (Fig 6H). Compared to crystal form I, the catalytic residue Asp412 was more distant from the γ-phosphate (∼8.5 Å) in these subunits. The nucleotide binding pattern is thus alternating in the ring, with the exception of the adjacent subunits G and A, which both harbor ATP and Mg^2+^. Interestingly, in two of the ADP-bound subunits (chains B and D), the tip of the αA-β2 loop, which cradles the nucleotide in the ATP-bound subunits, was remodeled and flipped away from ADP (compare Figs 6B and 6G). This conformational change might facilitate dissociation of ADP from the chaperonin.

### Cryo-EM structure of ɸEL in absence of nucleotide

To obtain the solution structure of apo-ɸEL, we next performed cryo-EM and single particle analysis. The raw micrographs and the particle 2D class averages suggested the presence of double-ring structures with approximate seven-fold rotational symmetry (S4A and S4B Figs). The particles appeared to have a strong tendency to associate via their apical domains (S4A Fig). Symmetry-free 3D class averaging indicated a tetradecameric double-ring structure with *C*2 symmetry (S4C Fig). Refinement of the particles resulted in a density map at 3.45 Å resolution (Fig 7A and S4D Fig), which allowed construction of an atomic model (Fig 7B and Table 2). The contacts between subunits were well defined in the density (Figs 7A and 7C). Similar to the crystal structures, the apical domains were the most mobile elements of the structure (S4E Fig). Apo-ɸEL has an open structure with a ring opening diameter at the apical domains of ∼65 Å (Fig 7A). This is in contrast to the previously reported 9 Å cryo-EM map of apo-ɸEL (EMD-6494) in which the ring opening is reduced to ∼27 Å due to rearrangement of the apical domains [10]. Overall each ring of the apo-ɸEL structure reported here is similar to the ɸEL heptamer of crystal form II (r.m.s.d. Cα positions 2.4 Å, versus 4.2 Å with crystal form I) in that the subunits alternate between conformations II (chains B, D, F) and III (chains A., C, E) and one subunit (chain G) is in conformation I (S5A-D Figs). Similar conformations of alternating subunits were recently observed by cryo-EM for the apo-state of the chaperonin of Pseudomonas phage OBP [50]. In all subunits of ɸEL the tip of the αA-β2 loop in the equatorial domain is moved away from the nucleotide binding site as in chains B and D of crystal form II (Fig 6G). Other nucleotide binding elements such as the P-loop (residues 79–86) and the β17-β18 hairpin (residues 503–512) are also moved outward (∼1.5 Å) in the apo structure. These local conformational changes might facilitate ATP-binding.

**Fig 7.**
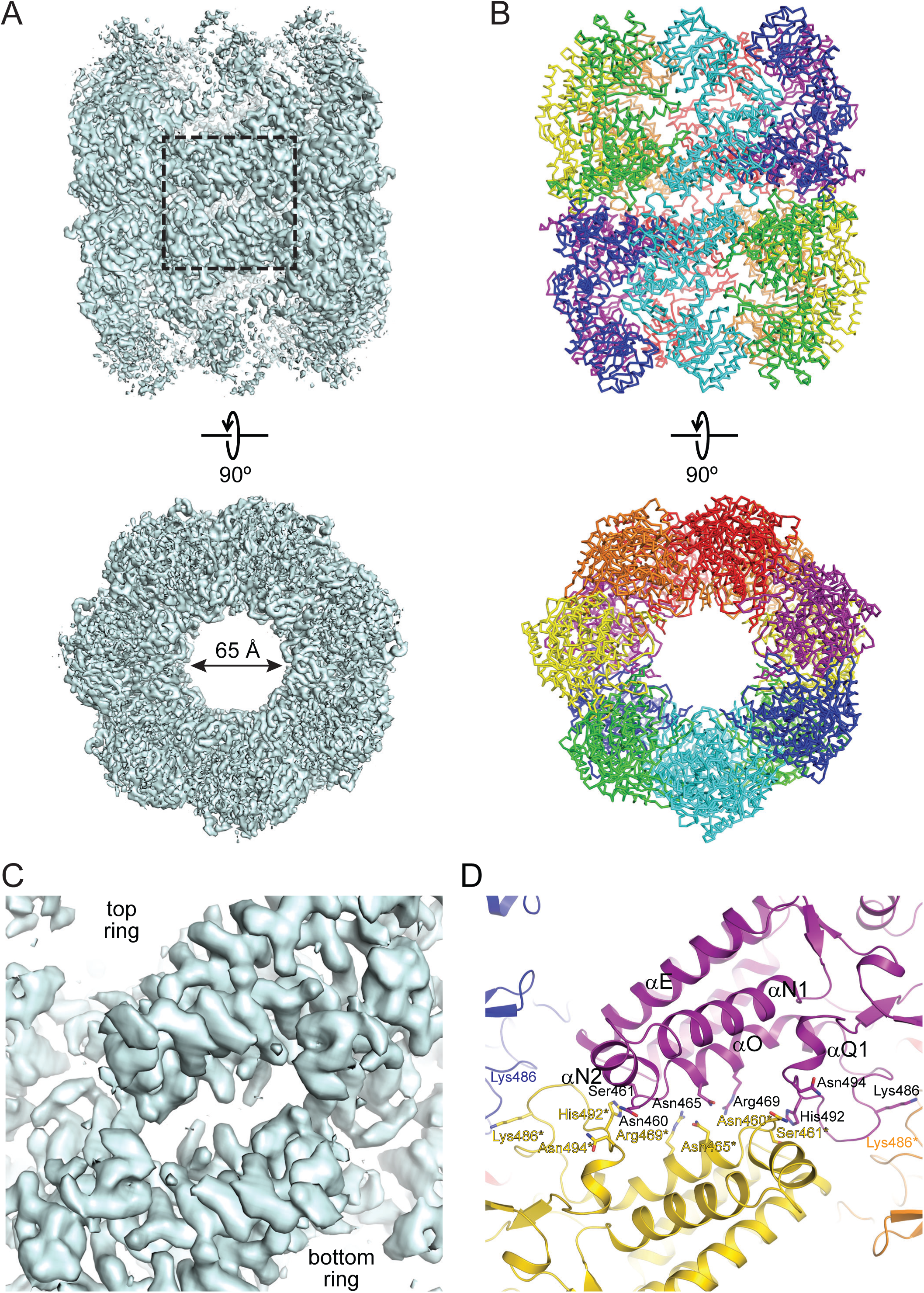
Cryo-EM structure of apo-ɸEL. (A) Electron density map of apo-ɸEL at 3.45 Å resolution. Perpendicular views of an isocontour surface at 4.8σ is shown. The top view is along the two-fold symmetry axis. The box highlights the inter-subunit interface at the equator of the complex. (B) Atomic model of apo-ɸEL. Backbone traces of the subunits are shown. Symmetry-equivalent subunits are shown in the same color. (C, D) Zoom on the symmetrical inter-subunit interface at the equator of the complex. Panel C shows the electron density. Panel D the model in ribbon representation. Contact sidechains are indicated and shown in stick representation (Glu466 is hidden behind the αO helix ribbon in this projection). Secondary structure elements participating in the interactions are indicated.

**Table 2.** Cryo-EM data collection and model statistics.

| ϕEL complex | Apo | ADP | ATPγS |
| --- | --- | --- | --- |
| EMDB number | 10528 | 10529 | 10530 |
| PDB code | 6TMV | 6TMW | 6TMX |
| **Data Collection** |  |  |  |
| Magnification | 81,000 | 73,000 | 73,000 |
| Voltage (kV) | 300 | 200 | 200 |
| Electron exposure (e<sup>-</sup> Å<sup>-2</sup>) | 78 | 43 | 42 |
| Defocus range (μm) | -1.5 to -3.0 | -1.5 to -4.0 | -1.5 to -3.5 |
| Pixel size (Å) | 1.090 | 1.997 | 1.997 |
| Symmetry imposed | *C*2 | *D*7 | *D*7 |
| Auto-picked features | 690,973 | 429,456 | 283,285 |
| Initial particle number | 356,175 | 408,474 | 137,262 |
| Final particle number | 178,107 | 170,957 | 52,885 |
| Map resolution (Å) | 3.45 | 5.9 | 5.8 |
| Map sharpening B factor (Å<sup>2</sup>) | -122 | -200 | -200 |
| **Model** |  |  |  |
| Resolution range (Å) | 209 – 3.45 | 192 – 5.9 | 192 – 5.8 |
| Average Fourier shell correlation | 0.824 | 0.925 | 0.875 |
| No. atoms |  |  |  |
| Protein | 58106 | 58058 | 58226 |
| Ligands | – | 378 | 462 |
| Average *B*-factors |  |  |  |
| Protein (Å<sup>2</sup>) | 209 | 286 | 298 |
| Ligands (Å<sup>2</sup>) | – | 120 | 150 |
| R.m.s. deviations |  |  |  |
| Bond length (Å) | 0.007 | 0.010 | 0.008 |
| Bond angles (°) | 1.131 | 1.433 | 1.114 |
| Ramachandran plot |  |  |  |
| Favored (%) | 96.1 | 94.2 | 96.1 |
| Allowed (%) | 3.9 | 5.6 | 3.9 |
| Outliers (%) | 0 | 0.2 | 0 |

Similar to group II chaperonins and unlike the staggered subunit contacts in group I, the ring-ring interface of apo-ɸEL is formed by two-fold symmetrical contacts between subunits. This results in the burial of 489–537 Å^2^ of accessible surface area (Figs 7C and 7D). For comparison, the buried surface area between subunits within heptamer rings is 1218-1401 Å^2^. The inter-ring contacts involve residues 492–494 in one subunit forming van-der-Waals contacts with residues 460* and 461* of the subunit in the opposite ring (Fig 7D). The negative dipole at the C-terminal end of helix αN2 is positioned close to His492*. Residues Asn465 and Asn465* form a symmetrical contact; Glu466 makes polar contacts with Arg469* and the backbone at Gly462*. Moreover, there is a symmetrical van-der-Waals contact with the neighboring subunit in the opposite ring, involving residues Lys486 (Fig 7D). None of these contacts are observed in group I and II chaperonins due to the high divergence of sequences.

### Cryo-EM structures of ɸEL in presence of ADP or ATPγS

To further elucidate the allosteric cycle of ɸEL, we determined the solution structure of ɸEL in the presence of ADP by single-particle cryo-EM. ɸEL formed double-ring structures with seven-fold rotational symmetry (S6A and S6B Figs). Symmetry-free 3D class averaging indicated an open double-ring structure with nearly uniform rings, and thus *D*7 symmetry was applied, resulting in a refined electron density map at 5.9 Å resolution (Fig 8A and S6C Fig). Note that the resolution is simply limited by the experimental setup. No particle class with closed single rings was found, in contrast to a previous report showing that ɸEL•ADP (EMD-6493) forms single rings resembling a hollow sphere [10] (S6B Fig). We used the domain structures from the 3.54 Å crystal structure to rigid-body fit and refine a pseudo-atomic molecular model of the ADP-bound ɸEL double-ring (Fig 8A and Table 2). In all the subunits, the αK-αL helical hairpin in the apical domain contacts the equatorial domain and the intermediate domain is in the inward-rotated conformation (subunit conformation I). The ring-ring interface was similar to that of apo-ɸEL (r.m.s.d. 1.4 Å for Cα atoms of all 14 equatorial domains) (Fig 8B).

**Fig 8.**
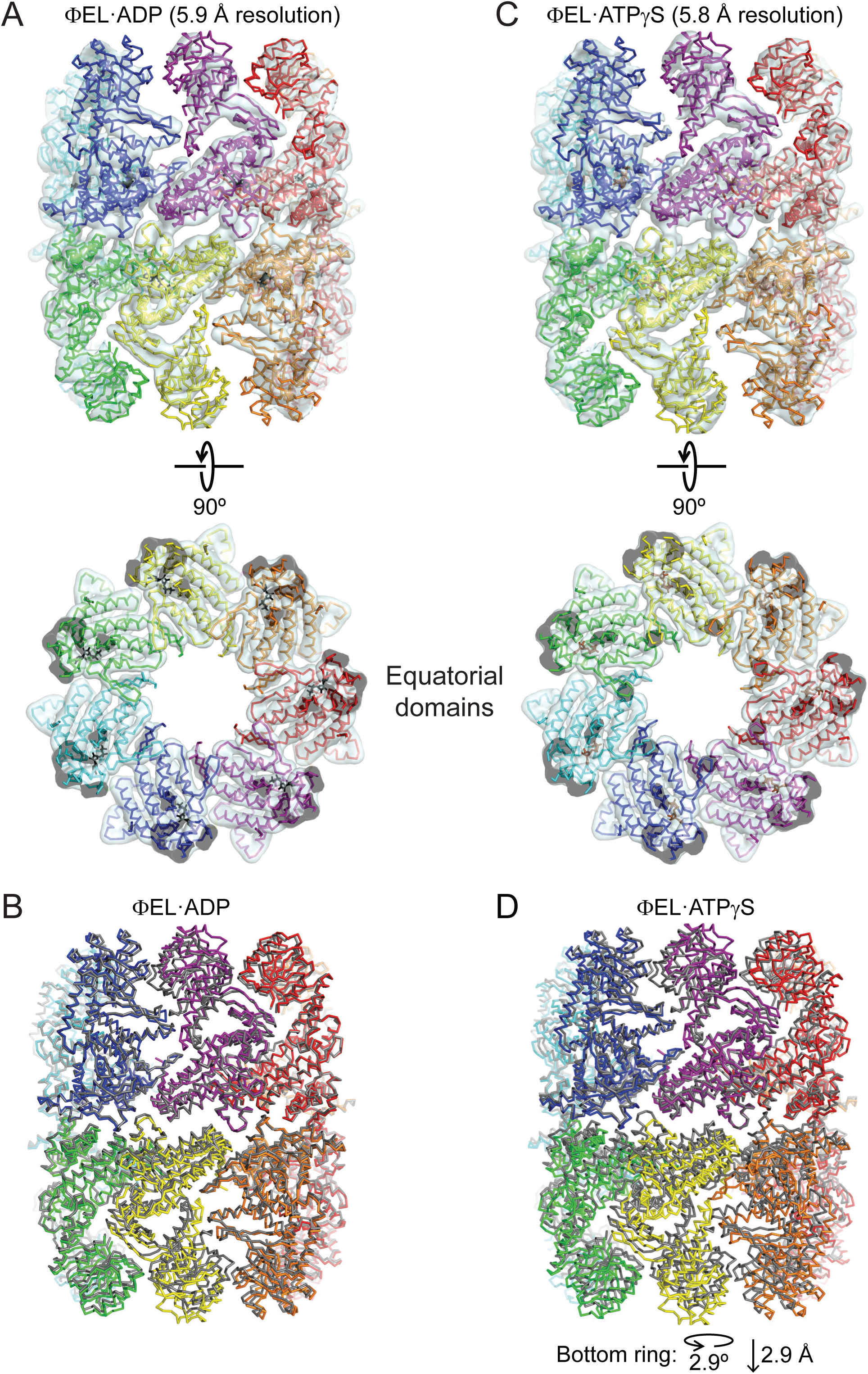
CryoEM structures of ɸEL in complex with ADP or ATPγS. (A) Superposition of the electron density map and the structural model of the ɸEL•ADP complex. The pseudo-atomic model is shown as backbone trace, with the subunits colored individually. ADP is shown in stick representation. The electron density map is shown as an isocontour surface at 4.5 σ. Two perpendicular views are shown. In the bottom view, only the equatorial domain section is shown to demonstrate the quality of the fit. (B) Superposition of the ɸEL•ADP complex with apo-ɸEL, showing the unaltered ring-ring interface. Backbone traces are shown. The ɸEL•ADP complex is shown in rainbow colors, and apo-ɸEL in grey. (C) Superposition of the electron density map and the structural model of the ɸEL•ATPγS complex. The representation is the same as in panel A. D) Superposition of the ɸEL•ATPγS complex with apo-ɸEL, showing the changes at the ring-ring interface. Backbone traces are shown. The ɸEL•ADP complex is shown in rainbow colors, and ɸEL•ATPγS in grey.

To mimic the conformational changes upon ATP-binding in solution, we also determined the cryo-EM structure of ɸEL in the presence of the slowly hydrolyzing ATP analog, ATPγS. Double-ring structures were again observed (S6D and S6E Figs) and symmetry-free 3D class averaging indicated nearly uniform rings. Applying *D*7 symmetry resulted in a refined electron density map at 5.8 Å resolution (Fig 8C and S6F Fig). The subunits assumed conformation I and were slightly tilted outward compared to ɸEL•ADP (Fig 8D). Interestingly, we find a rotation at the ring-ring interface of 2.9° with a vertical displacement of 2.9 Å (Fig 8D). This re-orientation would break the contacts between Asn465 and Glu466 with Asn465* and Arg469*/Gly462*, respectively (Fig 7D). Moreover, the symmetrical van-der-Waals contact at the Lys486 residues can no longer form. Thus, binding of ATPγS weakens the ring-ring interface and this effect may be more pronounced in the presence of ATP, consistent with the finding of mainly single-rings in solution (Fig 1C).

### Comparison of ɸEL with GroEL

Compared to GroEL of *E. coli*, ɸEL has numerous sequence insertions and deletions (S7 Fig). Only the nucleotide binding pocket is highly conserved. The apical domains of ɸEL differ in orientation from the apical domains of GroEL in both the open and GroES bound states (Figs 9A-C), but share the same secondary structure topology, i.e. the β-sandwich core structure, α-helices αH, αI and αJ at the surface and the helical hairpin formed by helices αK and αL (Figs 9D and 9E). However, the length of helices αH, αJ and αK and of the connecting surface loops differ substantially, resulting in a re-orientation of helices and re-modelling of long surface loops, such as residues 191–215 and 306–331, respectively (Fig 9D, boxed areas). The groove between helices αH and αI, which forms the binding site for substrate and GroES in GroEL, is more narrow and less deep in ɸEL. Accommodation of a helix or a β-hairpin element from SP would require structural remodeling of this site in ɸEL. In crystal form I, the putative SP binding site is solvent exposed, consistent with the ability of ɸEL to bind SP in the presence of ATP (Fig 4A). In contrast, in the apo-ɸEL (and crystal form II), helices αH and αI of chains A, C and E are partially occluded by the respective adjacent apical domain. The contact between the tip of the αK-αL helical hairpin to the equatorial domain in ɸEL is absent in GroEL, and the αK-αL connection is elongated and re-modelled in ɸEL (Figs 9A-C). The intermediate domains of GroEL and ɸEL are quite similar (Figs 9F and 9G), consistent with a conserved function in coupling domain movements with changes of the nucleotide status of the equatorial domain.

**Fig 9.**
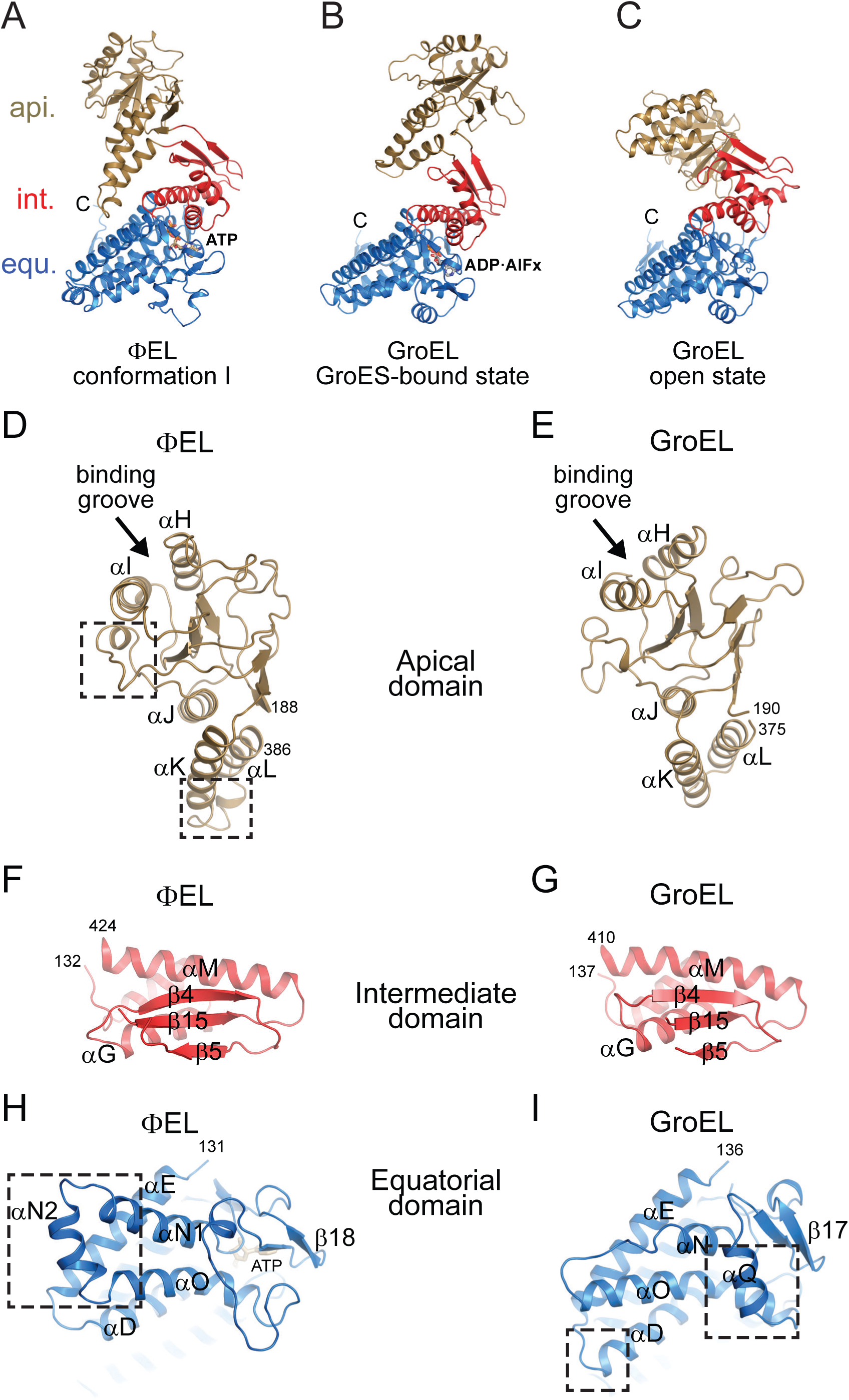
Comparison of the structures of ɸEL and GroEL. (A) Subunit structure of ɸEL (conformation I). The representation is the same as in Fig 6. (B, C) Structures of GroEL subunits from the *cis* (B) and *trans* rings (C) of the symmetric GroEL:GroES_2_ complex (PDB 1PCQ) [51]. The same representation style as in panel A. (D, E) Structures of the apical domains in ɸEL (D) and GroEL (E). Ribbon representations are shown. Remodeled regions in ɸEL discussed in the text are highlighted in boxes. Chain termini and α-helices are indicated. Arrows point to the proposed substrate binding site in group I chaperonins. For GroEL, the PDB dataset 1XCK (apo, open state) [52] was used. (F, G) Structures of the intermediate domains in ΦEL (F) and GroEL (G). Chain termini and selected secondary structure elements are indicated. (H, I) Structures of the equatorial domains in ɸEL (H) and GroEL (I). Ribbon representations are shown. A view from the ring-ring interface is shown. Large insertions in the respective structure involved in inter-ring contacts are highlighted in boxes.

The equatorial domains of ɸEL and GroEL differ mainly at the ring-ring interface contacts, with ɸEL making 1:1 subunit interactions and GroEL a 1:2 staggered interaction (Figs 9H and 9I). Helix αD is shortened in ɸEL by one turn compared to GroEL and helix αN is elongated by two turns (αN1), followed by insertion of a short helix, αN2. Helix αQ in GroEL (residues 462–471) is replaced by a loop connection in ɸEL (residues 479–503). This loop exhibits structural plasticity in the crystal structures of ɸEL single-ring.

Inter-domain salt bridges within subunits (equatorial domain D83 to apical domain K327) and between subunits (intermediate domain R197 to apical domain E386), which are important in allosteric regulation of GroEL [38], are not conserved in ɸEL.

## Discussion

Our structural and functional analysis of the chaperonin ɸEL from the bacteriophage EL of *P. aeruginosa* revealed that the protein is ATPase active and functions in aggregation prevention of denatured proteins. ɸEL forms tetradecameric double ring complexes, which dissociate into a population of single rings in the presence of ATP and at physiological salt concentration. In contrast, the recently observed dissociation of GroEL into single rings occurs only transiently during the reaction cycle [4]. The nucleotide bound single-ring complexes in the crystal structures of ɸEL closely resembled the individual rings of the double-ring complexes analyzed by cryo-EM. We could not confirm the existence of a sphere-like single-ring structure proposed to function in SP encapsulation [10]. Our functional data rather suggests that the chaperone mechanism of ɸEL is encapsulation independent, and represents an evolutionary precursor of the more complex encapsulation mechanisms used by the group I and II chaperonins [2, 5].

Based on our structural and functional analysis, we propose the following hypothetical model for the chaperonin cycle of ɸEL in SP binding and release, coupled to transitions between single- and double-ring complexes. Unlike group I and II chaperonins, the ability of ɸEL to prevent protein aggregation, as tested with Rho, as a model SP, was strictly nucleotide-dependent. In the apo-state ɸEL is a double ring with six of the seven subunits per ring assuming alternating states (conformations II and III) and one subunit in conformation I (Fig 10). The putative SP binding sites in the apical domains of subunits A, C and E are partially occluded. This form acquires competence in SP-binding upon binding of ATP, whereupon the apical domains of all seven subunits per ring are shifted to the same state (conformation I) and the equatorial domains are poised for ATP hydrolysis (Fig 10). ATP-binding weakens the ring-ring interface, resulting in single-ring formation at physiological salt concentration. SPs with a lower requirement for the availability of binding sites on adjacent subunits may also bind to the apo-state of ɸEL. ATP hydrolysis appears to occur in two stages: in the first stage, ATP is hydrolyzed in three alternating subunits, resulting in a ring conformation closely similar to the apo-state. This step may weaken the interaction with bound SP, perhaps allowing partial folding of some SPs. In the second stage, the remaining four subunits hydrolyze their ATP, followed by ADP dissociation generating the double-ring apo-state. This step presumably results in complete SP release for folding, as the apo-state is not SP-binding competent. Folding failed in the case of Rho, presumably because rebinding to ɸEL is faster than folding, consistent with the dependence of this slow folding protein on the encapsulation mechanism provided by GroEL/ES. Our structural analysis did not reveal an encapsulating state for ɸEL. Moreover, ɸEL did not functionally interact with GroES and does not contain a built-in-lid extension in its apical domains.

**Fig 10.**
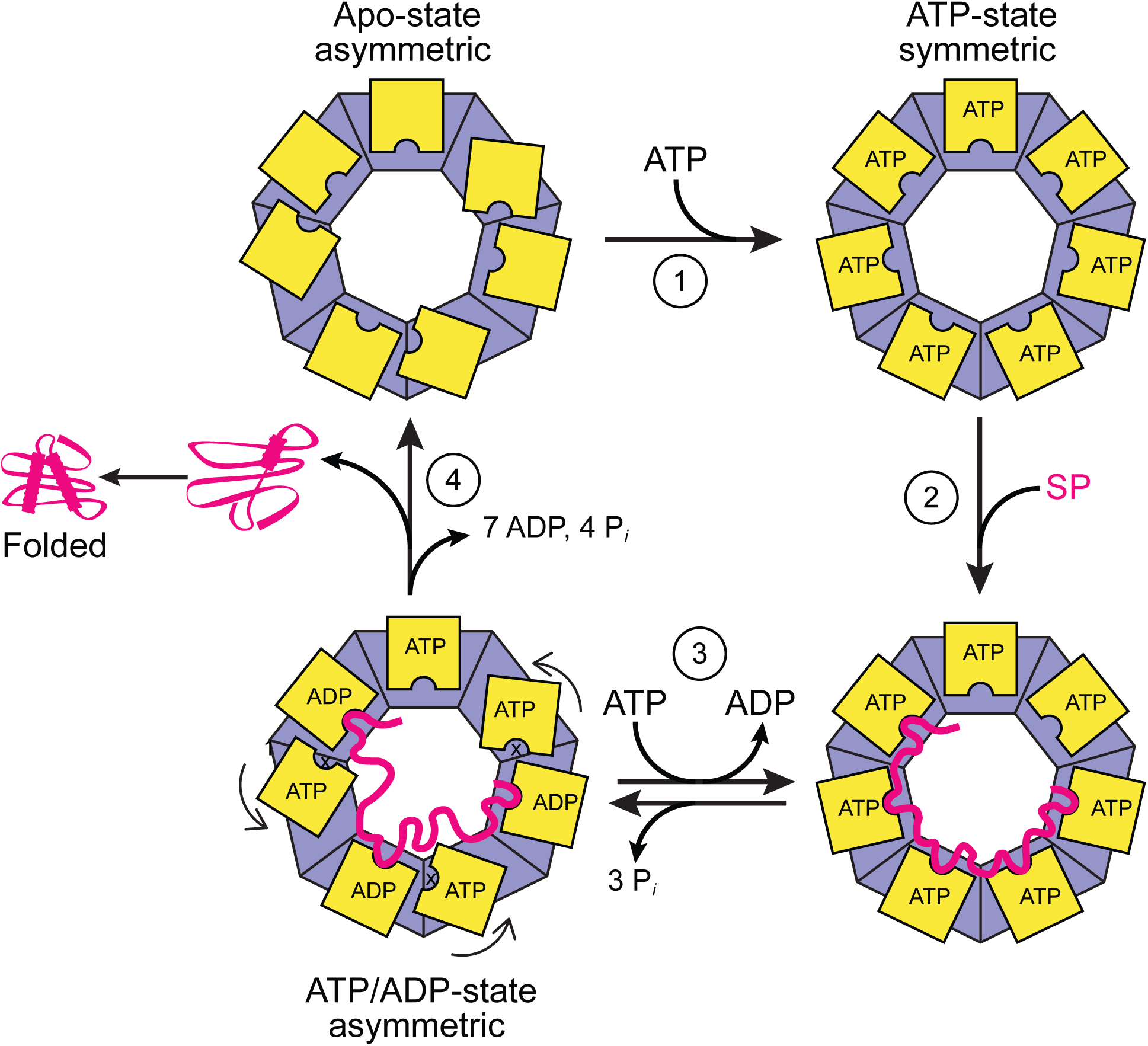
Hypothetical model for the ATPase and substrate interaction cycle of ɸEL. ɸEL is shown schematically as a top view with the apical domains in yellow. The double-ring apo-state with asymmetric apical domain orientation has only low affinity for SP. It is converted by ATP binding (1) to the single-ring characterized by symmetric apical domain topology and high binding affinity for SP (2). ATP hydrolysis in three alternating subunits (3) may result in partial SP release, with ATP hydrolysis in the remaining four subunits (4) completing SP release and folding.

Only some bacteriophage genomes encode their own chaperonin, presumably to assist in the folding of an essential phage protein(s) that cannot utilize (or does so only partially) the host chaperonin system for folding [10, 53]. Alternatively, ɸEL may prevent the folding of a phage protein up to a point when its function is required. For example, the putative ɸEL substrate gp188 is a cell wall endolysin [9] that should only function once a large number of phage particles have been produced by the host cell. ɸEL may stabilize gp188 in a non-native, inactive state until ATP levels may become depleted at the peak of phage production, resulting in concerted release from ɸEL and activation. It is also possible that additional regulatory factors play a role in triggering gp188 release. Future investigations will have to test the feasibility of such scenarios.

## Acknowledgements

We thank Lidia Kurochkina from the Russian Academy of Sciences, Moscow, Russia for providing us with the expression plasmid for ɸEL. We thank the JSBG group at the ESRF in Grenoble, France and the staff at the MPIB crystallization facility for their excellent support. Expert assistance by Tillman Schäfer and Daniel Bollschweiler from the MPIB cryo-EM facility is gratefully acknowledged. This work was supported by a grant from the Deutsche Forschungsgemeinschaft (DFG) (SFB1035) to M.H-H. and F.U.H.

## Author Contributions

A.B., F.U.H. and M.H-H conceived and designed the experiments. A.B., S.S.P., N.W. and M.H-H performed experiments. A.B., S.S.P., H.W., N.W and M.H-H analyzed the data. A.B., F.U.H. and M.H-H wrote the manuscript with contributions by all other authors.

## Supporting information

**S1 Fig.**
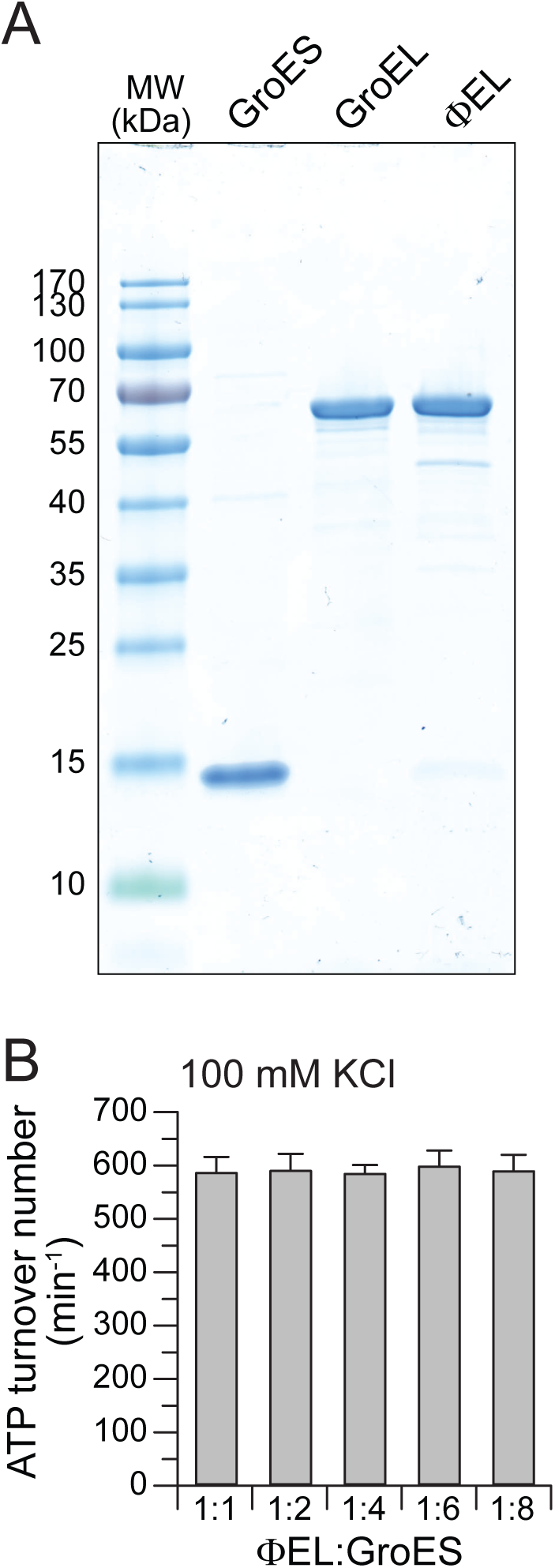
SDS-PAGE analysis of the ɸEL preparation. (A) A Coomassie Blue stained gel is shown. Purified GroEL and GroES were also analyzed. The positions of molecular weight markers are indicated. (B) Absence of effect of GroES on the ɸEL ATPase. ATPase measurements were performed with increasing molar excess of GroES heptamer over ɸEL tetradecamer, as indicated.

**S2 Fig.**
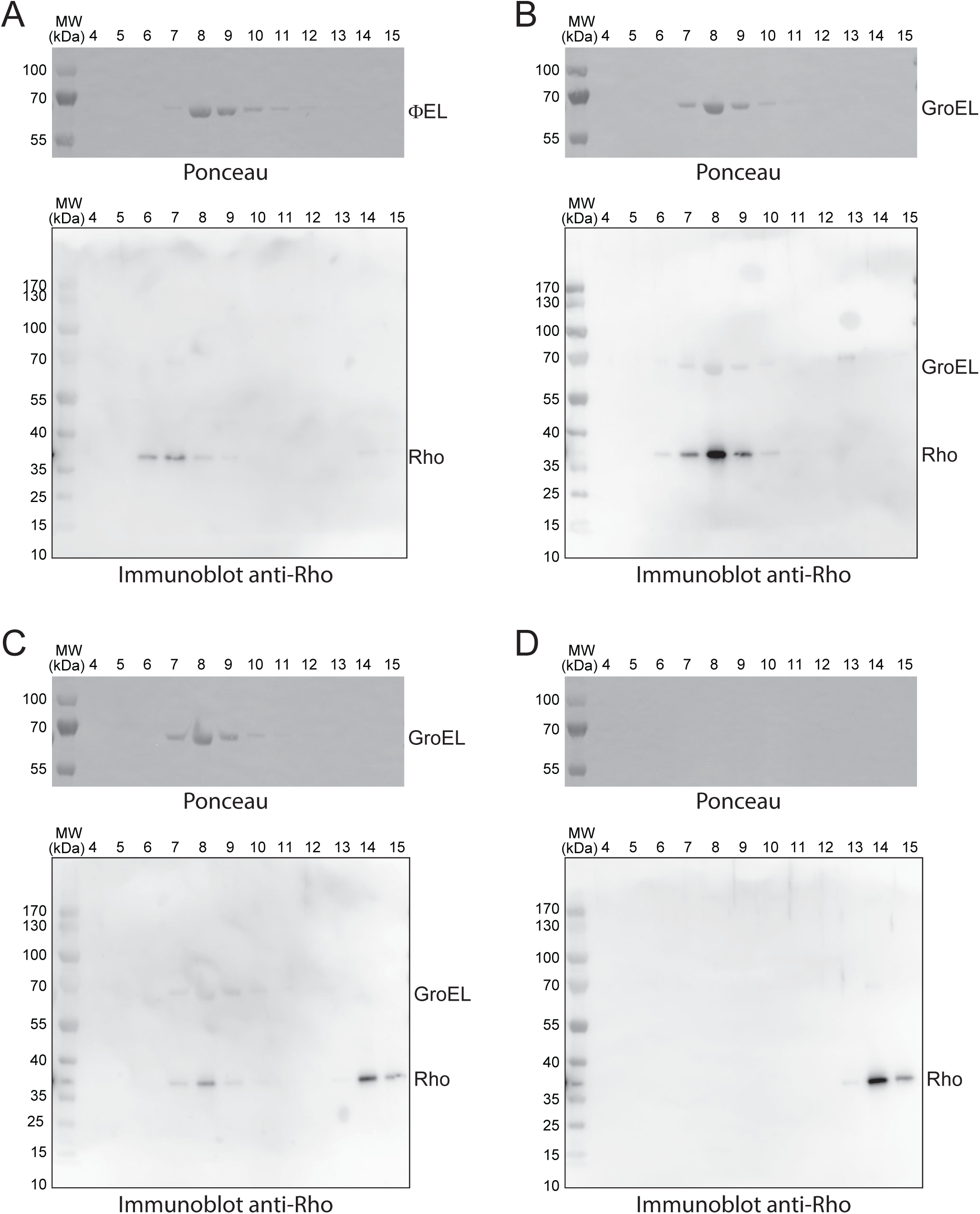
Size-exclusion chromatography ɸEL-rhodanese reactions. (A) ɸEL was incubated with denatured Rho in the presence of ATP for 30 min as in Fig 4C. The reaction was then analyzed by SEC with ATP in the column buffer, followed by Ponceau staining (top) and anti-Rho immunoblotting (bottom). (B and C) Rho folding reactions with GroEL/GroES in the absence of ATP (B) and in the presence of ATP (C). (D) Native Rho was analyzed as a control in the absence of chaperonin.

**S3 Fig.**
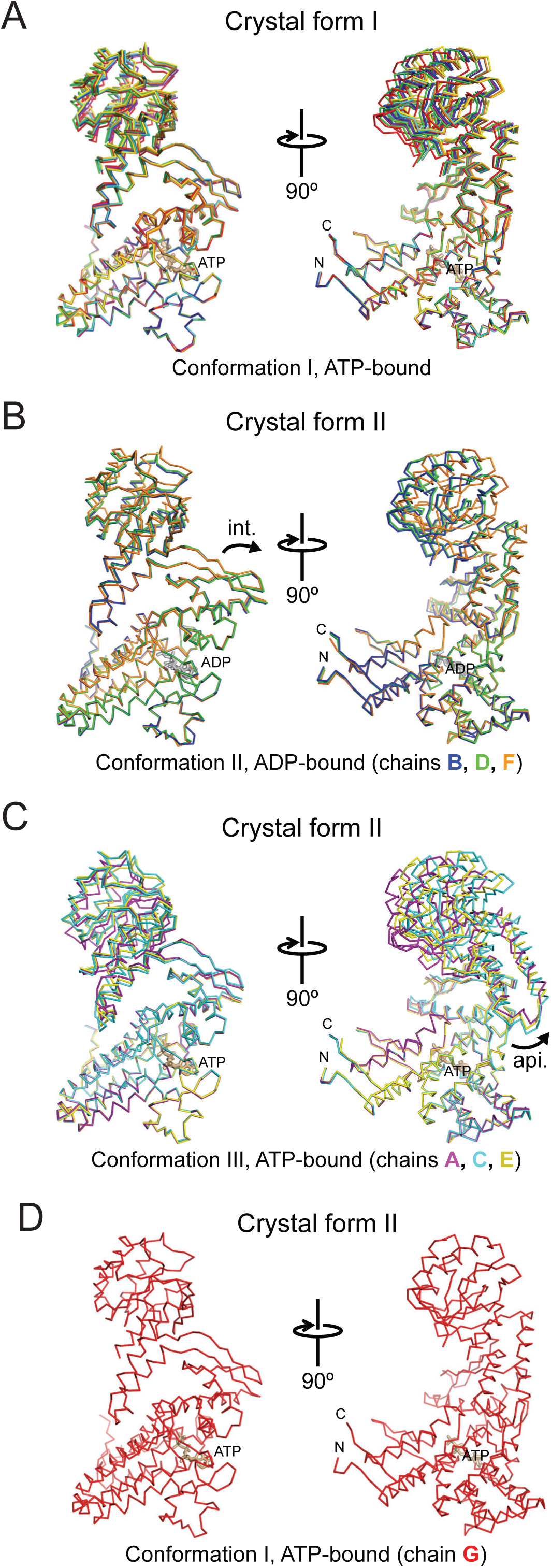
Subunit conformers found in ɸEL crystal structures. (A) Superposition of subunit conformers found in crystal form I. Perpendicular views of backbone traces are shown. The subunit colors are the same as in Fig 5. Bound ligand is shown in stick representation. N- and C-termini are indicated. The increased heterogeneity in the apical domain orientations (top) is caused by contacts in the crystal lattice. (B-D) Superposition of subunit conformers found in crystal form II. The subunits are grouped according to their conformation: Panels B, C and D show subunits in conformation II, III and I, respectively. Curved arrows indicate domain re-orientations relative to conformation I.

**S4 Fig.**
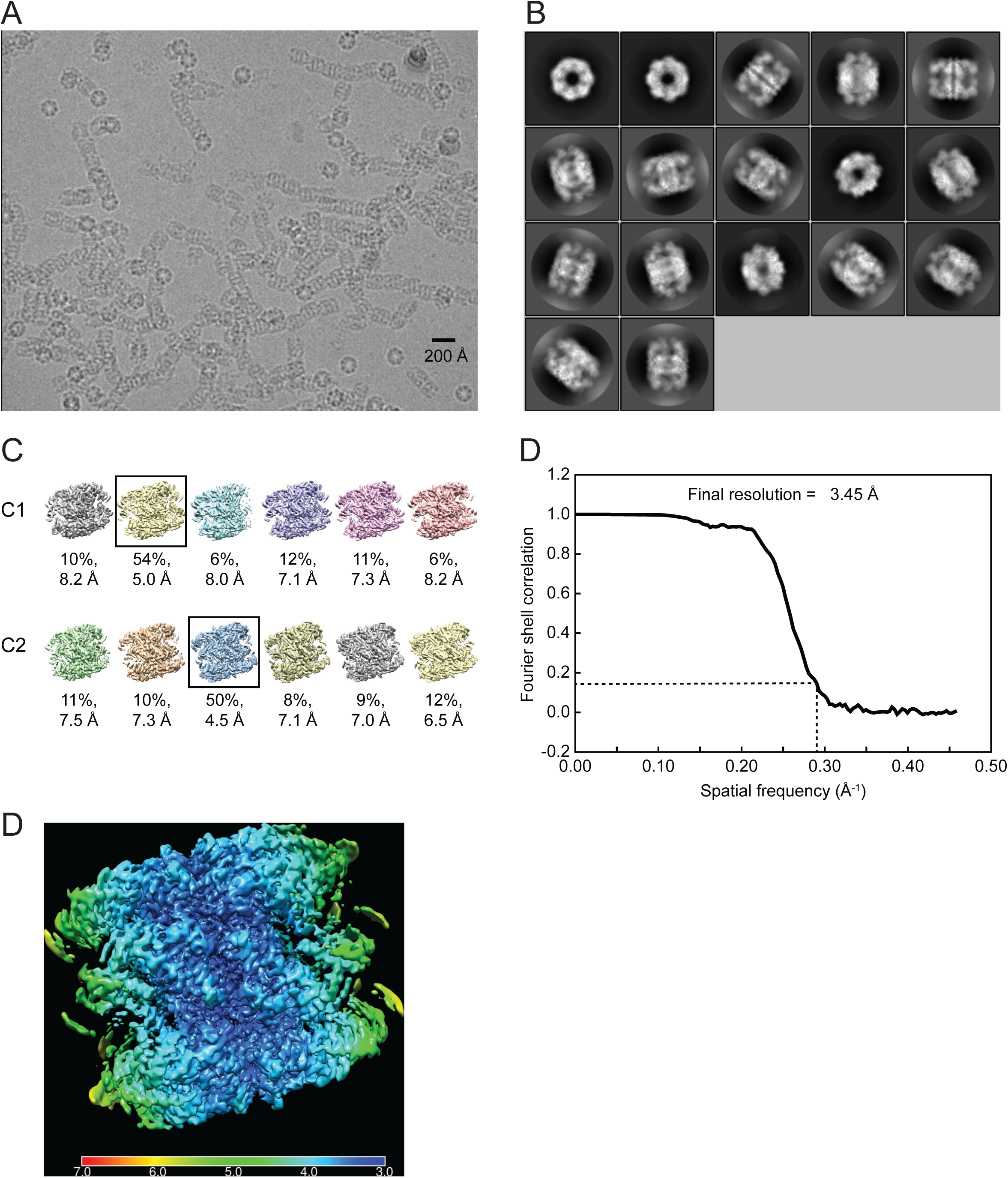
Cryo-EM analysis of apo-ɸEL. (A) Motion-corrected and dose-weighted raw micrograph of apo-ɸEL. For better contrast, the image was low pass filtered. The scale bar indicates 200 Å. (B) 2D class averages of apo-ɸEL particles. (C) Symmetry-free (top) and *C*2-symmetry averaged (bottom) 3D classes. The fraction of particles and the estimated resolution of the 3D classes are indicated. The refined classes are boxed. (D) Gold-standard FSC corrected curve of the final 3D reconstruction. The resolution was ∼3.45 Å at the FSC cutoff of 0.143. (E) Local resolution map for the apo-ɸEL structure. Local resolutions between 7 and 3 Å are represented as a rainbow color gradient from red to blue.

**S5 Fig.**
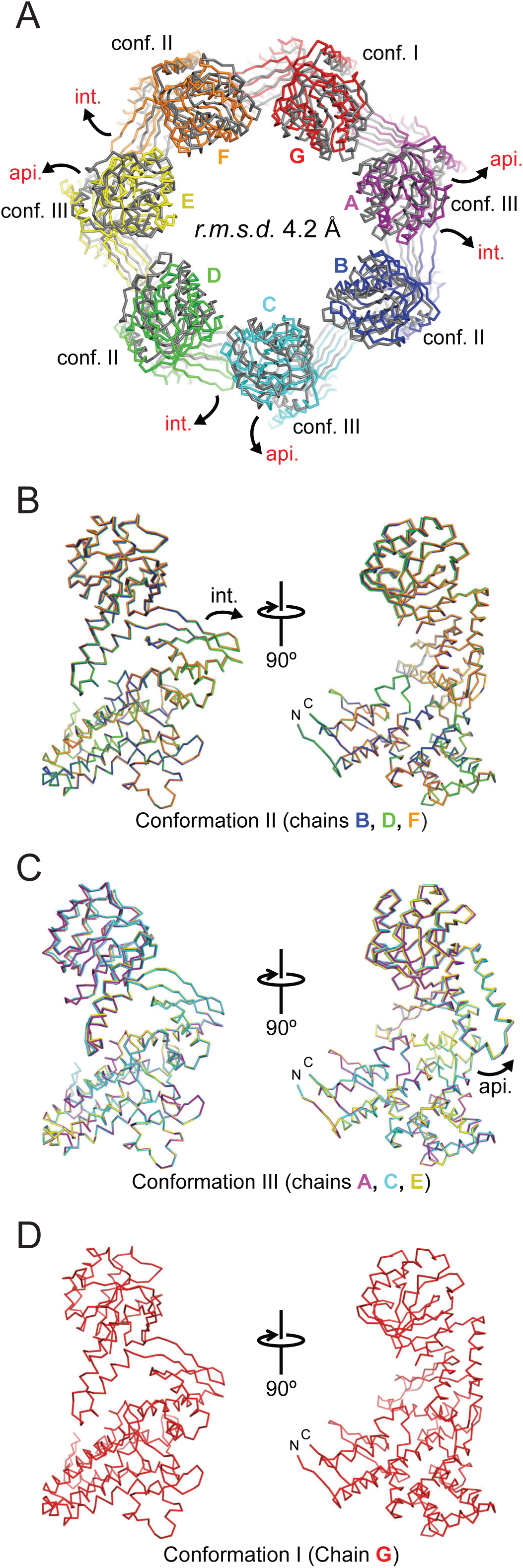
Conformational analysis of the apo-ɸEL structure. (A) Comparison of the subunits in the apo-ɸEL complex with crystal form I. Superposed backbone traces of only the apical and intermediate domains in a single ring are shown for clarity. The subunit colors of apo-ɸEL are the same as in Fig 7B. The crystal structure has grey color. Relative domain movements and conformation assignments are indicated. (B-D) Superposition of subunit conformers found in the cryo-EM structure of apo-ΦEL. The subunits are grouped according to their conformation: Panels B, C and D show subunits in conformation II, III and I, respectively. The subunit colors are the same as in Fig 7B.

**S6 Fig.**
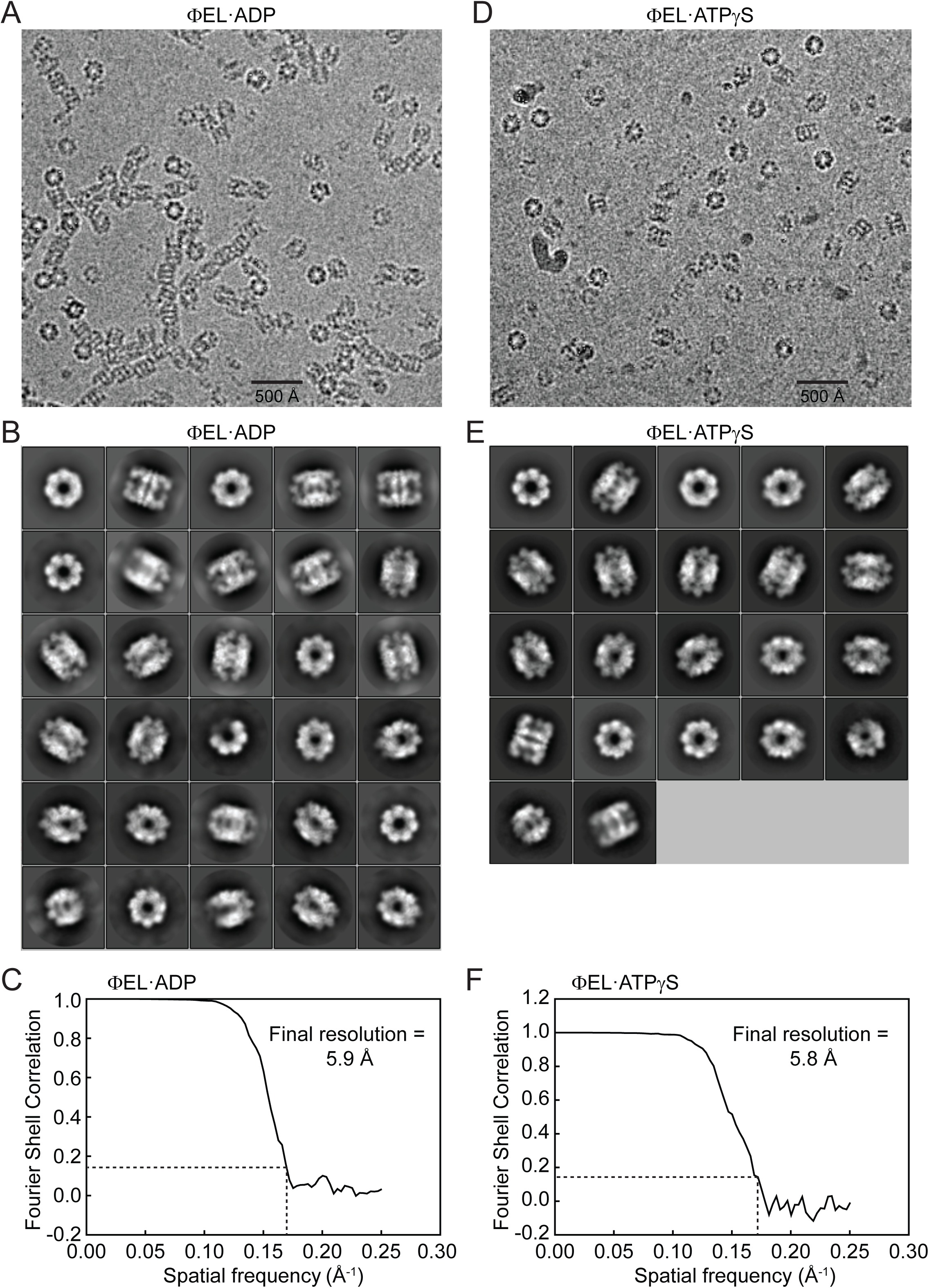
Cryo-EM analysis of the complexes of ɸEL with ADP and ATPγS. (A, D) Motion-corrected and dose-weighted raw micrographs of ɸEL•ADP (panel A) and ɸEL•ATPγS (D). The scale bar indicates 500 Å. (B, E) 2D class averages of ɸEL•ADP (B) and ɸEL•ATPγS (E) particles. C, F) Gold-standard FSC corrected curve (masked and B-factor sharpened) of the final 3D reconstructions of ɸEL•ADP (C) and ɸEL•ATPγS (F). The respective resolutions were ∼5.9 Å and ∼5.8 Å at the FSC cutoff of 0.143.

**S7 Fig.**
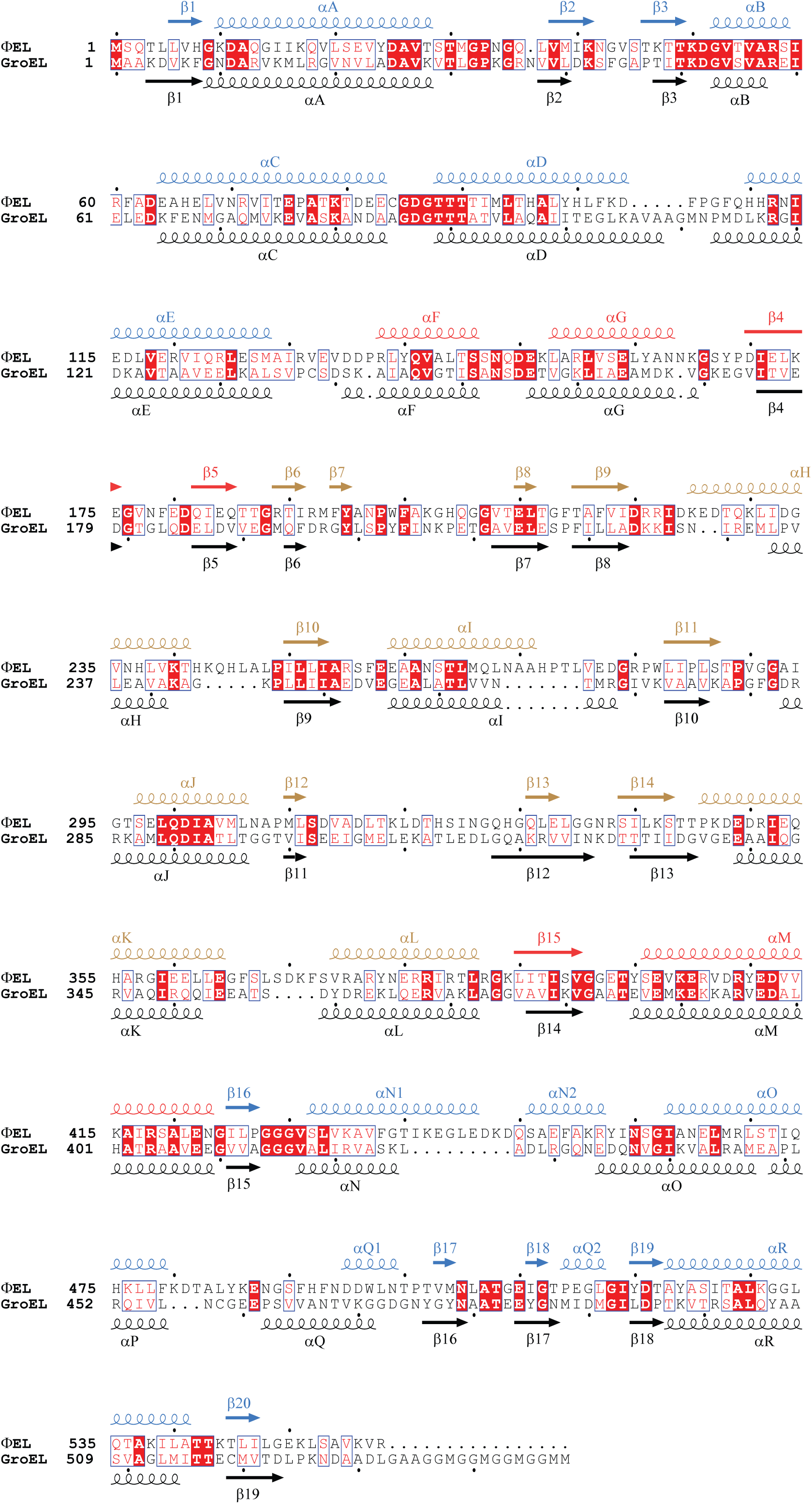
Sequence alignment of ɸEL and GroEL from *E. coli*. Secondary structure elements for ɸEL and GroEL from *E. coli* are indicated above and below the sequences, respectively. Similar residues are shown in red and identical residues in white on a red background. Blue frames indicate homologous regions. The Uniprot accession codes for the sequences are: Q2Z0T5, chaperonin from *Pseudomonas* phage EL; P0A6F5, GroEL from *E. coli*.

## References

1. Balchin D, Hayer-Hartl M, Hartl FU. *In vivo* aspects of protein folding and quality control. Science. 2016;353: aac4354. doi: 10.1126/science.aac4354.

2. Hayer-Hartl M, Bracher A, Hartl FU. The GroEL-GroES chaperonin machine: A nano-cage for protein folding. Trends Biochem Sci. 2016;41: 62–76. doi: 10.1016/j.tibs.2015.07.009.

3. Nielsen KL, Cowan NJ. A single ring is sufficient for productive chaperonin-mediated folding *in vivo*. Mol Cell. 1998;2: 93–9.

4. Yan X, Shi Q, Bracher A, Milicic G, Singh AK, Hartl FU, et al. GroEL ring separation and exchange in the chaperonin reaction. Cell. 2018;172: 605–17. doi: 10.1016/j.cell.2017.12.010.

5. Lopez T, Dalton K, Frydman J. The mechanism and function of group II chaperonins. J Mol Biol. 2015;427: 2919–30. doi: 10.1016/j.jmb.2015.04.013.

6. van der Vies SM, Gatenby AA, Georgopoulos C. Bacteriophage T4 encodes a co-chaperonin that can substitute for *Escherichia coli* GroES in protein folding. Nature. 1994;368: 654–6. doi: 10.1038/368654a0.

7. Marine RL, Nasko DJ, Wray J, Polson SW, Wommack KE. Novel chaperonins are prevalent in the virioplankton and demonstrate links to viral biology and ecology. ISME J. 2017;11: 2479–91. doi: 10.1038/ismej.2017.102.

8. Hertveldt K, Lavigne R, Pleteneva E, Sernova N, Kurochkina L, Korchevskii R, et al. Genome comparison of *Pseudomonas aeruginosa* large phages. J Mol Biol. 2005;354: 536–45. doi: 10.1016/j.jmb.2005.08.075.

9. Kurochkina LP, Semenyuk PI, Orlov VN, Robben J, Sykilinda NN, Mesyanzhinov VV. Expression and functional characterization of the first bacteriophage-encoded chaperonin. J Virol. 2012;86: 10103–11. doi: 10.1128/JVI.00940-12.

10. Molugu SK, Hildenbrand ZL, Morgan DG, Sherman MB, He L, Georgopoulos C, et al. Ring separation highlights the protein-folding mechanism used by the Phage EL-encoded chaperonin. Structure. 2016;24: 537–46. doi: 10.1016/j.str.2016.02.006.

11. Tafoya DA, Hildenbrand ZL, Herrera N, Molugu SK, Mesyanzhinov VV, Miroshnikov KA, et al. Enzymatic characterization of a lysin encoded by bacteriophage EL. Bacteriophage. 2013;3: e25449. doi: 10.4161/bact.25449.

12. Wyatt PJ. Light-scattering and the absolute characterization of macromolecules. Analytica Chimica Acta. 1993;272: 1–40. doi: 10.1016/0003-2670(93)80373-S.

13. Poso D, Clarke AR, Burston SG. A kinetic analysis of the nucleotide-induced allosteric transitions in a single-ring mutant of GroEL. J Mol Biol. 2004;338: 969–77. doi: 10.1016/j.jmb.2004.03.010.

14. Weber F, Hayer-Hartl M. Refolding of bovine mitochondrial rhodanese by chaperonins GroEL and GroES. Methods Mol Biol. 2000;140: 117–26. doi: 10.1385/1-59259-061-6:117.

15. Radaev S, Li S, Sun PD. A survey of protein-protein complex crystallizations. Acta Crystallogr D Biol Crystallogr. 2006;62: 605–12.

16. Radaev S, Sun PD. Crystallization of protein-protein complexes. J App Cryst. 2002;35: 674–6.

17. Kabsch W. XDS. Acta Crystallogr D Biol Crystallogr. 2010;66: 125–32. doi: 10.1107/S0907444909047337.

18. Evans P. Scaling and assessment of data quality. Acta Crystallogr D Biol Crystallogr. 2006;62: 72–82. doi: 10.1107/S0907444905036693.

19. Evans PR, Murshudov GN. How good are my data and what is the resolution? Acta Crystallogr D Biol Crystallogr. 2013;69: 1204–14. doi: 10.1107/S0907444913000061.

20. French G, Wilson K. On the treatment of negative intensity observations. Acta Cryst Sect A. 1978;34: 517–25. doi: 10.1107/S0567739478001114.

21. Potterton E, Briggs P, Turkenburg M, Dodson E. A graphical user interface to the CCP4 program suite. Acta Crystallogr D Biol Crystallogr. 2003;59: 1131–7.

22. Matthews BW. Solvent content of protein crystals. J Mol Biol. 1968;33: 491–7. doi: 10.1016/0022-2836(68)90205-2.

23. Vagin AA, Isupov MN. Spherically averaged phased translation function and its application to the search for molecules and fragments in electron-density maps. Acta Crystallogr D Biol Crystallogr. 2001;57: 1451–6.

24. Pettersen EF, Goddard TD, Huang CC, Couch GS, Greenblatt DM, Meng EC, et al. UCSF Chimera--a visualization system for exploratory research and analysis. J Comput Chem. 2004;25: 1605–12. doi: 10.1002/jcc.20084.

25. Terwilliger TC. Maximum-likelihood density modification. Acta Crystallogr D Biol Crystallogr. 2000;56: 965–72.

26. Emsley P, Cowtan K. Coot: model-building tools for molecular graphics. Acta Crystallogr D Biol Crystallogr. 2004;60: 2126–32.

27. Murshudov GN, Skubak P, Lebedev AA, Pannu NS, Steiner RA, Nicholls RA, et al. REFMAC5 for the refinement of macromolecular crystal structures. Acta Crystallogr D Biol Crystallogr. 2011;67: 355–67. doi: 10.1107/S0907444911001314.

28. Zheng SQ, Palovcak E, Armache JP, Verba KA, Cheng Y, Agard DA. MotionCor2: anisotropic correction of beam-induced motion for improved cryo-electron microscopy. Nat Methods. 2017;14: 331–2. doi: 10.1038/nmeth.4193.

29. Rohou A, Grigorieff N. CTFFIND4: Fast and accurate defocus estimation from electron micrographs. J Struct Biol. 2015;192: 216–21. doi: 10.1016/j.jsb.2015.08.008.

30. Scheres SH. RELION: implementation of a Bayesian approach to cryo-EM structure determination. J Struct Biol. 2012;180: 519–30. doi: 10.1016/j.jsb.2012.09.006.

31. Zivanov J, Nakane T, Forsberg BO, Kimanius D, Hagen WJ, Lindahl E, et al. New tools for automated high-resolution cryo-EM structure determination in RELION-3. Elife. 2018;7. doi: 10.7554/eLife.42166.

32. Schorb M, Haberbosch I, Hagen WJH, Schwab Y, Mastronarde DN. Software tools for automated transmission electron microscopy. Nat Methods. 2019;16: 471–7. doi: 10.1038/s41592-019-0396-9.

33. Mastronarde DN. Automated electron microscope tomography using robust prediction of specimen movements. J Struct Biol. 2005;152: 36–51. doi: 10.1016/j.jsb.2005.07.007.

34. Biyani N, Righetto RD, McLeod R, Caujolle-Bert D, Castano-Diez D, Goldie KN, et al. Focus: The interface between data collection and data processing in cryo-EM. J Struct Biol. 2017;198: 124–33. doi: 10.1016/j.jsb.2017.03.007.

35. Chen VB, Arendall WB, 3rd, Headd JJ, Keedy DA, Immormino RM, Kapral GJ, et al. MolProbity: all-atom structure validation for macromolecular crystallography. Acta Crystallogr D Biol Crystallogr. 2010;66: 12–21. doi: 10.1107/S0907444909042073.

36. Kleywegt GT, Jones TA. A super position. CCP4/ESF-EACBM Newsletter on Protein Crystallography. 1994;31: 9–14.

37. Gouet P, Courcelle E, Stuart DI, Metoz F. ESPript: multiple sequence alignments in PostScript. Bioinformatics. 1999;15: 305–8.

38. Gruber R, Horovitz A. Allosteric mechanisms in chaperonin machines. Chem Rev. 2016;116: 6588–606. doi: 10.1021/acs.chemrev.5b00556.

39. Langer T, Pfeifer G, Martin J, Baumeister W, Hartl FU. Chaperonin-mediated protein folding: GroES binds to one end of the GroEL cylinder, which accommodates the protein substrate within its central cavity. EMBO J. 1992;11: 4757–65.

40. Martin J, Geromanos S, Tempst P, Hartl FU. Identification of nucleotide-binding regions in the chaperonin proteins GroEL and GroES. Nature. 1993;366: 279–82. doi: 10.1038/366279a0.

41. Kawe M, Plückthun A. GroEL walks the fine line: the subtle balance of substrate and co-chaperonin binding by GroEL. A combinatorial investigation by design, selection and screening. J Mol Biol. 2006;357: 411–26. doi: 10.1016/j.jmb.2005.12.005.

42. Hayer-Hartl MK, Weber F, Hartl FU. Mechanism of chaperonin action: GroES binding and release can drive GroEL-mediated protein folding in the absence of ATP hydrolysis. EMBO J. 1996;15: 6111–21.

43. Tang YC, Chang HC, Roeben A, Wischnewski D, Wischnewski N, Kerner MJ, et al. Structural features of the GroEL-GroES nano-cage required for rapid folding of encapsulated protein. Cell. 2006;125: 903–14. doi: 10.1016/j.cell.2006.04.027.

44. Gupta AJ, Haldar S, Milicic G, Hartl FU, Hayer-Hartl M. Active cage mechanism of chaperonin-assisted protein folding demonstrated at single-molecule level. J Mol Biol. 2014;426: 2739–54. doi: 10.1016/j.jmb.2014.04.018.

45. Martin J, Langer T, Boteva R, Schramel A, Horwich AL, Hartl FU. Chaperonin-mediated protein folding at the surface of GroEL through a ‘molten globule’-like intermediate. Nature. 1991;352: 36–42. doi: 10.1038/352036a0.

46. Mayhew M, da Silva AC, Martin J, Erdjument-Bromage H, Tempst P, Hartl FU. Protein folding in the central cavity of the GroEL-GroES chaperonin complex. Nature. 1996;379: 420–6. doi: 10.1038/379420a0.

47. Weissman JS, Rye HS, Fenton WA, Beechem JM, Horwich AL. Characterization of the active intermediate of a GroEL-GroES-mediated protein folding reaction. Cell. 1996;84: 481–90. doi: 10.1016/s0092-8674(00)81293-3.

48. Brinker A, Pfeifer G, Kerner MJ, Naylor DJ, Hartl FU, Hayer-Hartl M. Dual function of protein confinement in chaperonin-assisted protein folding. Cell. 2001;107: 223–33. doi: 10.1016/s0092-8674(01)00517-7.

49. Hayward S, Berendsen HJ. Systematic analysis of domain motions in proteins from conformational change: new results on citrate synthase and T4 lysozyme. Proteins. 1998;30: 144–54.

50. Stanishneva-Konovalova TB, Semenyuk PI, Kurochkina LP, Pichkur EB, Vasilyev AL, Kovalchuk MV, et al. Cryo-EM reveals an asymmetry in a novel single-ring viral chaperonin. J Struct Biol. 2019:107439. doi: 10.1016/j.jsb.2019.107439.

51. Chaudhry C, Farr GW, Todd MJ, Rye HS, Brunger AT, Adams PD, et al. Role of the gamma-phosphate of ATP in triggering protein folding by GroEL-GroES: function, structure and energetics. EMBO J. 2003;22: 4877–87. doi: 10.1093/emboj/cdg477.

52. Bartolucci C, Lamba D, Grazulis S, Manakova E, Heumann H. Crystal structure of wild-type chaperonin GroEL. J Mol Biol. 2005;354: 940–51. doi: 10.1016/j.jmb.2005.09.096.

53. Semenyuk PI, Orlov VN, Kurochkina LP. Effect of chaperonin encoded by gene 146 on thermal aggregation of lytic proteins of bacteriophage EL *Pseudomonas aeruginosa*. Biochemistry 2015;80: 172–9. doi: 10.1134/S0006297915020042.

